# Sex-specific disease association and genetic architecture of retinal vascular traits

**DOI:** 10.1101/2025.07.16.665150

**Authors:** Leah Böttger, Dennis Bontempi, Olga Trofimova, Michael J Beyeler, Sacha Bors, Ilaria Iuliani, Ian Quintas, David Presby, Sven Bergmann

**Affiliations:** Department of Computational Biology, University of Lausanne, Lausanne, Switzerland; Swiss Institute of Bioinformatics, Lausanne, Switzerland; Department of Integrative Biomedical Sciences, University of Cape Town, Cape Town, South Africa

## Abstract

Sex as a biological variable (SABV) remains underemphasized in biomedical research, despite its well-established impact on disease prevalence, progression, and genetic architecture. Here, we investigated sex differences in retinal vascular phenotypes, which are emerging as non-invasive biomarkers for ocular, cardiovascular, and neurodegenerative diseases. We used both interaction and sex-stratified analyses in disease association and genome-wide association studies (GWAS), including the X chromosome. Our analyses revealed sex-specific differences in associations between retinal traits, cardiometabolic risk factors, and disease outcomes. Several phenotypes showed prognostic value for mortality and cardiovascular endpoints, predominantly in one sex, illustrating the limitations of sex-agnostic analyses. Genetic analyses revealed differences in genetic architectures, with females exhibiting higher heritability and a greater number of associated loci, as well as sex-specific trait-gene associations. Notably, we report the first genetic association on the X chromosome for retinal vascular phenotypes. These findings suggest sex-differential genetic and environmental contributions to microvascular morphology, possibly reflecting divergent biological pathways and risk profiles. Our study underscores the necessity of integrating SABV and sex-aware approaches in genetic and prognostic research to advance precision medicine and improve tailored risk stratification.

## Introduction

Decades of research have highlighted the important but complex role of sex as a biological variable (SABV) for understanding phenotypic diversity, disease prevalence, and risk (Gualtierotti, 2025). Since the 1990s, major funding agencies such as the US National Institutes of Health (NIH), the Canadian Institutes of Health Research (CIHR), and the European Commission (EC) have called for integrating sex and gender into biomedical research agendas (White *et al*., 2021). Yet, as detailed below, the implementation of these recommendations in research still lags behind.

Physiological sex or gender (for a definition, see **Suppl. Info 1**) differences extend far beyond reproductive traits. Research has shown significant disparities in disease susceptibility, clinical manifestation, progression, and mortality, e.g., in autoimmune disorders (Angum *et al*., 2020), neurodegenerative (Pinares-Garcia *et al*., 2018), and cardiovascular diseases (Appelman *et al*., 2015; Stanhewicz, Wenner and Stachenfeld, 2018; Vaura *et al*., 2022), as well as differences in a range of complex phenotypes. The mechanisms underlying these sex differences are multifaceted, involving hormonal, genetic, and environmental factors (Khramtsova, Davis and Stranger, 2019). For cardiovascular diseases in particular, the role of gonadal hormones, especially estrogens, has continuously been highlighted as a female protective factor (Stanhewicz, Wenner and Stachenfeld, 2018), with risk for women increasing after menopause (Ryczkowska *et al*., 2023).

The sex chromosomes play a crucial role in differences between the sexes, not only through determining the differentiation of gonads and expression of sex hormones but also through their interactions with autosomal genes and environmental factors (Bale and Epperson, 2017). Patterns of X chromosomal gene expression further impact phenotypic differentiation between the sexes due to different dosages, i.e., females having two copies whereas males only have one, parental imprinting, and the presence, absence or skewing of X-chromosome inactivation (Arnold, Chen and Itoh, 2012; Khramtsova, Davis and Stranger, 2019). Nevertheless, the X chromosome remains underexplored, even after repeated calls for its inclusion (Wise, Gyi and Manolio, 2013). Only 25% of studies in the NHGRI-EBI GWAS catalog report results related to X-linked variation (Sun *et al*., 2023b). More recently, large-scale biobank studies have revealed sex differences in genetic architectures for a range of complex traits and diseases. Whereas effect differences between females and males were relatively small, they were widespread across the genome and showed diverging patterns of heritability, cross-trait correlations, as well as tissue-specific gene expression (Kassam *et al*., 2019; Oliva *et al*., 2020; Bernabeu *et al*., 2021). Yet, the majority of large GWAS studies continue to report results based only on mixed samples, potentially overlooking sex-specific or differentiated effects.

Despite the ample evidence of their importance, sex differences in physiological and genomic research are still widely ignored (Arnegard *et al*., 2020; Merone *et al*., 2022), resulting in incomplete or biased understandings of biological mechanisms, which continue to contribute to growing health disparities. In this study, we aim to explore sex differences in disease associations and genetic architectures in the retinal vasculature, an easily accessible biomarker which is under frequent investigation as a potentially powerful screening tool for a range of ocular, cardiovascular, and neurodegenerative disorders (Ikram *et al*., 2013; Chandra *et al*., 2019; Allon *et al*., 2021; Snyder *et al*., 2021; Tomasoni *et al*., 2023; Ortín Vela *et al*., 2024). Leveraging the vascular phenotypes from Ortín Vela et al. (2024) extracted from UK Biobank (UKB) color fundus photographs (CFPs), we examined how SABV influenced association results through phenotypic or genetic interaction and investigated sex-specific disease association and genetic architectures. Our focus on the retinal vasculature, a non-invasive biomarker for systemic, cardiovascular, and neurodegenerative disorders, is motivated by its clinical relevance in the context of precision medicine, underscoring the importance of sex-specific approaches in diagnosis, treatment, and disease risk stratification.

## Results

### Sex differences in phenotypic traits

Our first aim was to investigate whether there are any sex-specific differences in the image-derived phenotypes (IDPs). Before statistical analyses, we applied quality control measures to account for potential sex-specific confounders: we excluded individuals with sex chromosome aneuploidies (n=104) and discrepancies between genetic and reported sex (n=33). We visually compared the distributions of the 17 IDPs and quantified their mean differences in units of standard deviations (Cohen’s *d*, where *d* = (*F̅* − *M̅*) / *std*(*pooled*)).

Trait distributions between the sexes were largely overlapping for all IDPs with Cohen’s *d* ranging from −0.143 to 0.220 (**Suppl. Fig. 1**). We found the largest observed differences in the arterial (A) temporal angle (*d* = −0.143), A median diameter (*d* = 0.161), and the arterial over venous (A/V) ratio vascular density (*d* = 0.220). Despite small differences in distributions, all traits showed significant mean differences except venous (V) central retinal equivalent (eq).

### Sex differences in disease association

To study differences in disease or risk factor association of our IDPs, we used multiple linear regression for continuous variables (risk factors) and multiple logistic regression for binary variables (disease states). We used two complementary approaches: 1. an interaction analysis (IDP x sex) to investigate if associations differed by sex, and 2. a sex-stratified approach to independently estimate effect sizes for each sex and compare them using Z-statistics (Clogg, Petkova and Haritou, 1995). For more details, please see the methods.

In the interaction analyses, we observed significant IDP × sex interaction effects for 10 out of 17 IDPs across several risk factors (**Suppl. Fig. 3a**). We found the largest number of sex-differing associations for bifurcations (8) and A vascular density (8), mainly for blood markers (i.e., low-density lipoprotein (LDL), triglycerides) and blood pressure but also for smoking. These findings suggest that the relationship between several of our IDPs differs significantly between the sexes, but they do not provide sex-specific effect size estimates adjusted independently for all covariates.

To characterize differences in disease associations between females and males, we estimated effect sizes independently for each cohort. In concordance with Ortin-Vela et al. (2024), we found significant associations for multiple trait-disease and risk factor pairs, even for reduced sex-specific sample sizes (**Suppl. Fig. 3b-c**). While effect sizes and directions were generally consistent between the sexes, differences were mostly observed in the level of statistical significance, with more associations reaching significance in males than in females. For example, effects highly significant in males but not in females included A central retinal eq for glaucoma, A standard deviation of (short std) diameter for death, and V vascular density for diabetes. Conversely, associations significant in females but not in males included V tortuosity for diabetes, A vascular density for stroke, and A central retinal eq for hypertension. Notable exceptions with a difference in sign as well as significance where males showed a significant association but not females were found for five trait-disease/ -risk pairs: A median diameter and atherosclerosis, V median diameter and CVD, A central retinal eq and LDL, as well bifurcations and A vascular density and smoking pack-years.

We found effect differences to be significant for several diseases or risk factors across multiple traits. These were most frequent in A vascular density (11), V tortuosity (10), and bifurcations (7), and more pronounced in the risk factors. We found highly significant differences where females showed more positive or less negative effect sizes for the pairs BMI - V tortuosity, pulse rate - A/V ratio vascular density, and smoking pack-years - bifurcations/ A vascular density with the same pattern in males for systolic blood pressure - A vascular density, as well as pulse rate - V std diameter/ V central retinal eq (**Fig. 1**).

**Figure 1.**
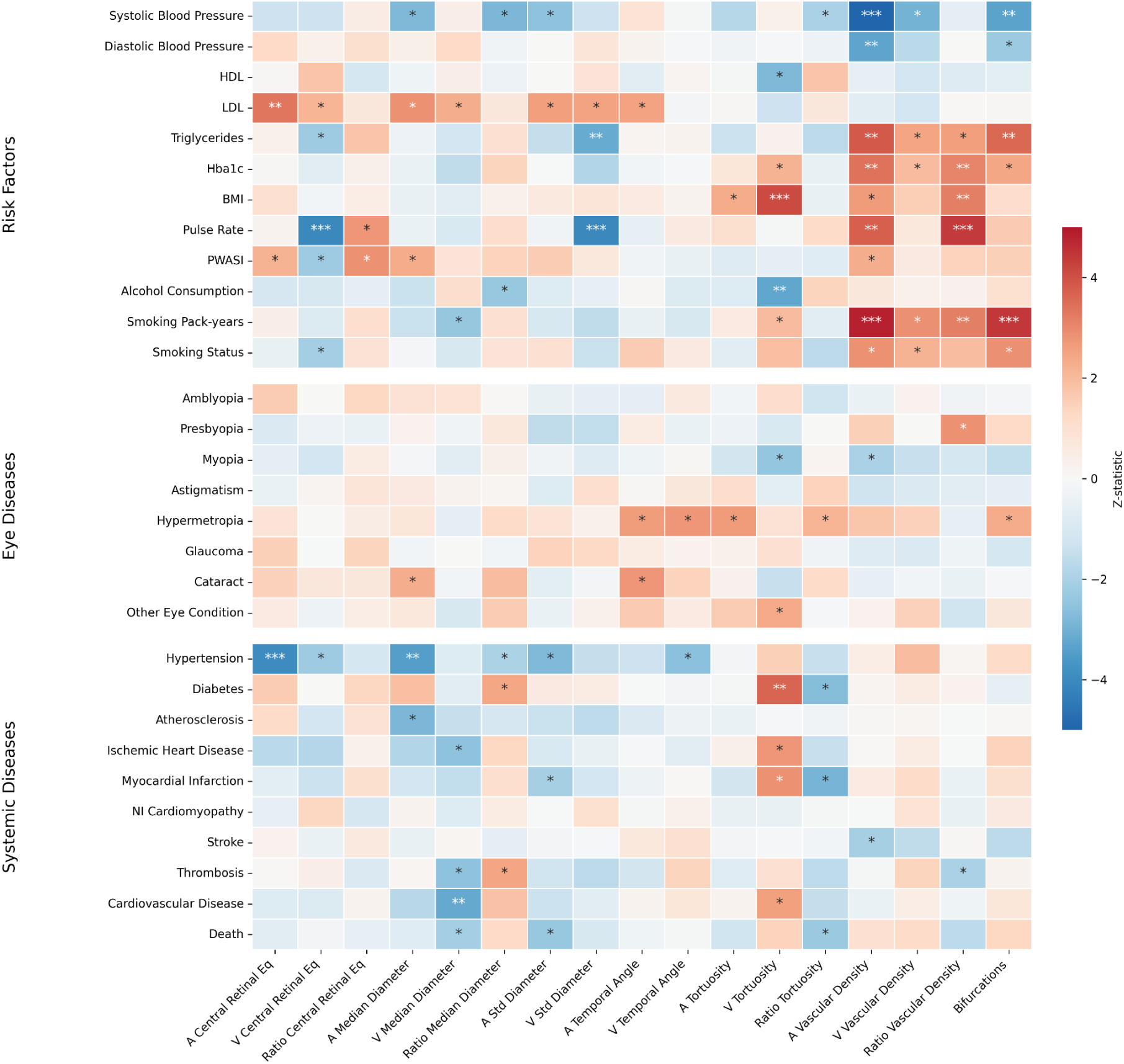
Sex differences in risk factor and disease association for all 17 IDPs. Estimation of sex differences in regression coefficients using Z-tests shows that several phenotypic traits show significantly differing association with risk factors or diseases between females and males. A positive Z value indicates a more positive association in females, negative Z values indicate a more positive association in males. Asterisks indicate the level of statistical significance: ∗ = p < 0.05, ∗∗ = p < 0.05/ N_IDPs_, where N_IDPs_= 17, and ∗∗∗ = p < 0.05/ N_IDPs_ x N_Diseases_, where N_Diseases_ = 31 was considered the number of all dependent variables used in the linear and logistic regressions. BMI: body mass index; HDL: high-density lipoprotein; LDL: low-density lipoprotein; NI: Nonischemic; PWASI: Pulse Wave Arterial Stiffness Index; A: arterial; V: venous; Eq: equivalent; Std: standard deviation; ratio: A over V ratio.

We further compared our complementary approaches to each other by examining the concordance of their effect estimates (**Suppl. Fig. 3d**). Overall, results were largely consistent between both methods. Whereas the interaction analysis yielded more nominally significant trait-disease or -risk associations, some sex differences, most prominently for the risk factors but also for diseases such as diabetes, myopia, or stroke, were only detected in the Z-statistic. We observed a discrepancy with high-magnitude interaction estimates, mainly for the A and V vascular density and bifurcations in relation to LDL and systolic blood pressure (SBP), which were not corroborated, at least not to the same extent, by the Z-test.

### Survival Analysis

To investigate sex-specific prognostic power of the IDPs, we evaluated their association with cardiovascular outcomes and death (all-cause mortality) using Cox Proportional Hazard (PH) models. For every IDP-outcome pair, we fit three independent models - one for the whole cohort, one for females only, and one for males only - adjusting for major cardiovascular risk factors (see methods). UKB fields used to define all covariates and endpoints can be found in **Suppl. Tab. 2.1-2.3**, average follow-up per endpoint in **Suppl. Tab. 2.5**, and distributions of the date of event for different endpoints in **Suppl. Fig. 4a-c**.

In the sex-stratified survival analysis, we observed that the effect sizes of the association between various IDPs and endpoints, measured by Hazard Ratios (HRs), differed between males, females, and the combined cohort (**Fig. 2**). In 6 cases, IDPs were significantly associated with the endpoints across all three models: in two instances (A vascular density for CVD/IHD) effect sizes were similar, whereas A and V vascular density, as well as bifurcations, showed lower HRs for all-cause mortality (death) in males and bifurcations exhibited a lower HR for females for stroke. We identified 20 cases where one sex exclusively contributed to any associations observed in the overall analysis (14 cases for males and 6 for females), indicating that the specific trait was independently prognostic only within a subset of the cohort. For example, V median diameter was associated with all-cause mortality in the combined analysis (HR=1.05, 95%CI 1.02–1.08, p<0.01). However, stratified analyses revealed that the trait was prognostic in males (HR=1.07 (1.03–1.11), p<0.001) but not in females (HR=1.01 (0.965–1.06), p=0.683). Most notably, we found 7 cases where no significant association was observed in the combined cohort, with the opposite happening in one of the sex-stratified models (5 for males and 2 for females). A/V ratio vascular density was not associated with all-cause mortality in the overall analysis (HR=1.03, 95%CI 1.00–1.06, p=0.0565) or the female cohort (HR=0.988 (0.944–1.03), p=0.586), but it was in males (HR=1.06 (1.02–1.10), p<0.01). This was also the case in A temporal angle and CVD death (Comb: HR=1.09 (0.999–1.19), p=0.053; F: HR=0.962 (0.925–0.999), p<0.05); M: HR=1.18 (1.06–1.31), p<0.01); A/V ratio median diameter and CVD (Comb: HR=0.986 (0.964–1.01), p=0.218; F: HR=1.01 (0.974–1.05), p=0.586; M: HR=0.969 (0.941–0.997), p<0.05); V central retinal eq (Comb: HR=1.03 (0.987–1.08), p=0.162; F: HR=0.982 (0.903–1.07), p=0.660); M: HR=1.06 (1.00–1.12), p<0.05) and A vascular density, and MI events (Comb: HR=0.952 (0.904–1.00), p=0.0661; F: HR=0.977 (0.889–1.08), p=0.621; M: HR=0.938 (0.881–0.999), p<0.05). Conversely, A/V ratio tortuosity was not associated with all-cause mortality in the combined analysis (HR=0.983, 95%CI 0.955–1.01, p=0.230) or the male cohort (HR=1.01 (0.971–1.05), p=0.691), but it was in females (HR=0.948 (0.906–0.992), p<0.05). A central retinal eq and CV events were further only significantly associated in females (Comb: HR=0.987 (0.964–1.01), p=0.265; F: HR=0.962 (0.925–0.999), p<0.05; M: HR=1.00 (0.975–1.03), p=0.979).

**Figure 2.**
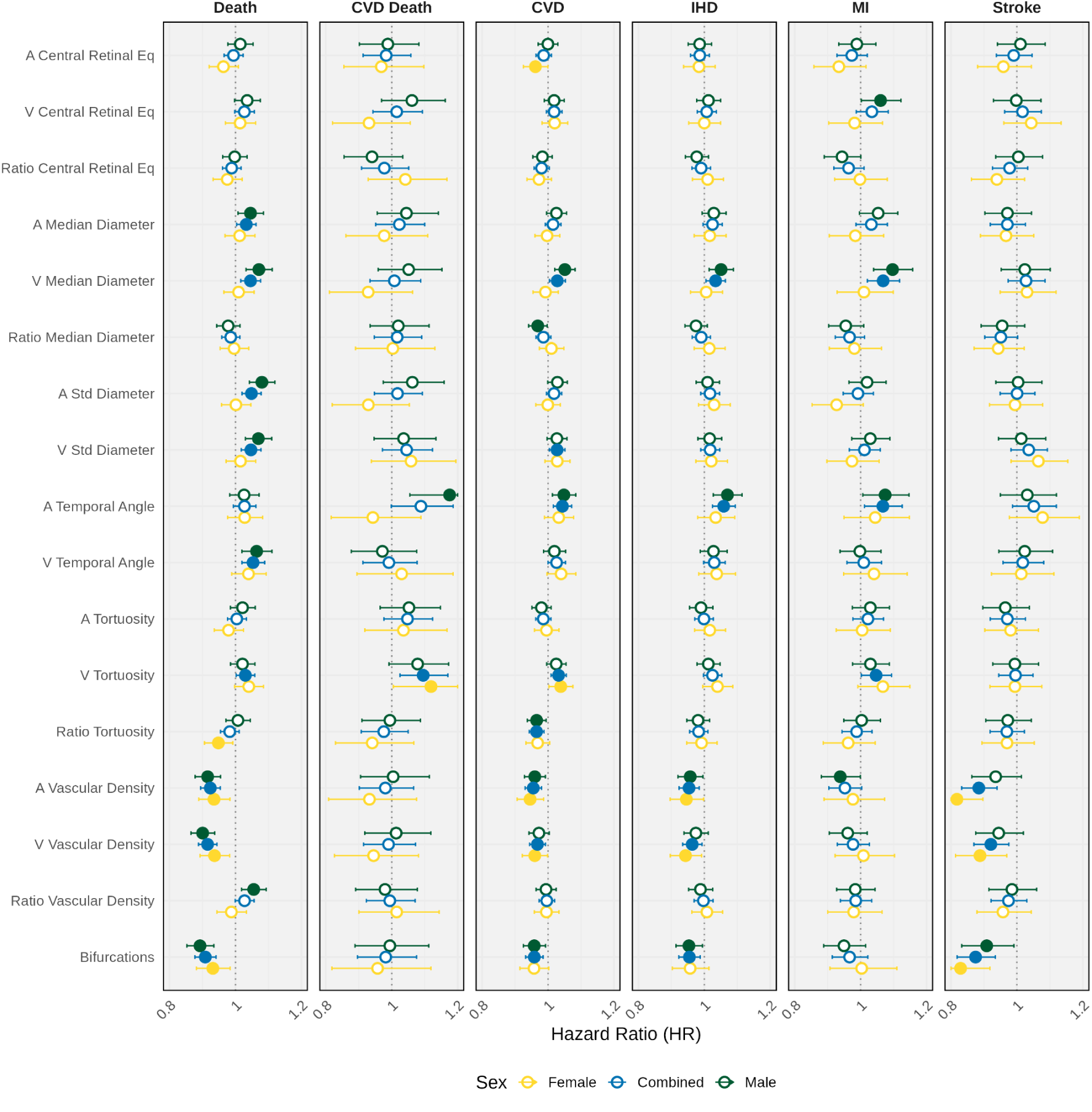
Hazard Ratios and 95% Confidence Intervals for the 17 IDPs and the selected endpoints. Sex-stratified Cox proportional hazards models between IDPs and cardiovascular outcomes or all-cause mortality. Full circles represent significant hazard ratios (p<0.05). CVD: cardiovascular (event); IHD: ischaemic heart disease; MI: myocardial infarction.

### Sex-specific Genetic Architecture of the retinal microvasculature

We conducted Genome-wide association studies (GWAS) for the 17 IDPs in both sex-stratified as well as the combined cohort, including the X chromosome in our analysis pipeline. We used a similar approach as for the disease association: we first assessed potential sex-differential effects using SNP x Sex interaction analysis and subsequently performed sex-stratified GWAS independently using the same set of covariates, further quantifying significant effect differences using Z-tests. Results from the sex-stratified analyses were then meta-analyzed to estimate pooled effect sizes and investigate potential effect heterogeneity as well as compare to the combined sample analysis. We used PascalX (Krefl, Brandulas Cammarata and Bergmann, 2023) for gene enrichment analysis for the autosomes and the X chromosome (see Methods).

The SNP x Sex interaction results were mainly insignificant (**Suppl. Fig. 5i-j**), indicating that for the majority of loci, the genetic effects on retinal microvasculature traits did not differ significantly by sex. The only exception was A/V ratio vascular density, where a relatively large amount of SNPs (n = 1615) distributed across all chromosomes, with a notable peak on Chr 15 (p < 1e-30), reached genome-wide significance. Five traits, i.e., A tortuosity, A vascular density, A/V ratio vascular density, V temporal angle, and V tortuosity, showed medium to high levels of genomic inflation (1.20 < λ < 1.64) possibly due to sex-specific trait variance, thus violating the assumption of homoscedasticity, or due to interaction-specific model misspecifications, such as unaccounted confounding variables or differential covariate effects between sexes.

Concordant with the findings of Ortin-Vela et al. (2024), all IDPs showed a polygenic architecture, even in the smaller female and male cohorts (**Suppl. Fig. 5a-f**, see **Fig. 3a** for aggregated results of all IDPs). The inclusion of the X chromosome into the GWAS yielded 5 new independent variants in the combined analysis, none of which have been previously associated with a quantitative or disease trait. These significant associations were found for rs5916123 (A central retinal equivalent), rs5961501 (A tortuosity), and rs626840 (A temporal angle). For V temporal angle, two SNPs on the X chromosome were significantly associated: rs113876691 and rs626840, the latter being an intron variant of the gene EFNB1. rs68126241 (A std diameter and A/V ratio tortuosity) was the only SNP on the X chromosome that was also significant in the smaller male cohort, whereas none reached significance in females.

**Figure 3.**
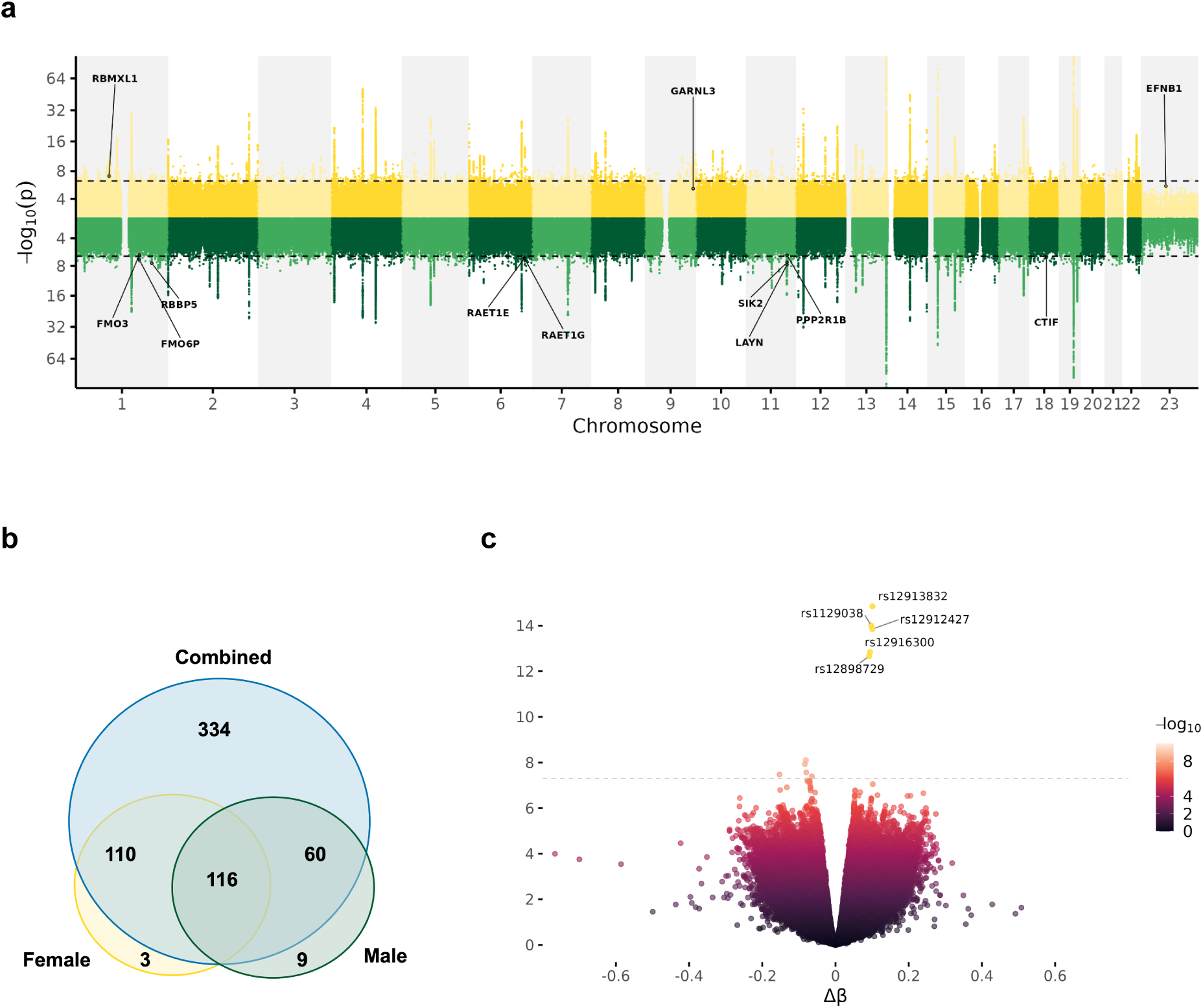
SNP- and Gene-wise sex differences. a. Miami plot for Female and Male GWAS results showing-log10 p significance values across the 17 traits on a log2 transformed scale. Genes with sex-specific associations are indicated. 23 denotes the X chromosome. b. Gene-wise overlap between male and female cohorts c. Volcano plot of SNP-wise effect differences. SNPs significant in the individual female or male GWAS and significantly different according to the Z-tests (both p < 5e-8) are colored with their respective color (Female = gold, Male = green).

In the combined sample, a total of 25 976 SNPs were associated with one or more of the retinal traits while the number of associated loci in the individual sex-stratified analyses was substantially lower (9 267 in females, 7 802 in males). We identified 903 sex-specific SNPs (496 in females, 407 in males) that did not reach nominal genome-wide significance in the combined sample (**Fig. 3b**). Interestingly, our meta-analyses (**Suppl. Fig. 5g-h**) showed fewer significant SNPs (n = 20 215) than the combined analyses (n = 25 976). Of these, 724 were unique to the meta-analyses, 89 were shared exclusively with the male cohort, and 88 with the female cohort. We found only 4 496 SNPs to be significant in all four cohorts, and, on average, more overlap between the combined or meta-analyses and the female cohort (**Suppl. Fig. 5l**).

Comparing all IDP combined GWAS results (representative of the retinal vasculature), we found that females, on average, had more significant variants per chromosome, most prominently on chromosomes 15 (F – M = 661), 17 (F – M = 451), and 1 (F – M = 450), while males only had remarkably more significant SNPs on chromosome 8 (M – F = 947). 10 loci showed significant differences in their beta coefficients (Δβ = β_*F*_ − β_*M*_) of which 5 were genome-wide significant (**Fig. 3c**) in the female cohort for A/V ratio vascular density, showing sex-specific trait-by-genotype distributions (**Suppl. Fig. 6a**). All five SNPs are located within the gene HERC2 on chromosome 15 and are in moderate to perfect linkage disequilibrium (LD) (r² ranging from 0.644 [rs12898729–rs12912427] to 1 [rs1129038–rs12913832], **Suppl. Info. 3**). To further investigate how these SNPs might drive phenotypic differences, we adjusted A/V ratio vascular density for the sex-specific SNP effect in each cohort [*ratio vascular density*_*SNP*_ = *ratio vascular density* − (β_*SNP*_ * *genotype*)], and recomputed their distributions. This adjustment largely reduced phenotypic differences between the females and males, with Cohen’s *d* decreasing substantially (Cohen’s *d_SNP_*ranging from 0.02 to 0.04 depending on SNP, **Suppl. Fig. 6b).**

PascalX gene scoring results, similar to our SNP results, revealed a substantially lower number of genes associated with each sex-stratified sample (Comb = 620, F = 229, M = 185). 116 genes were shared between all cohorts (**Fig. 3b**) whereas three were found to be unique to females and nine to males (annotated for their approximate location in **Fig. 3a**). For females, these were *RBMXL1* (chromosome 1) and *GARNL3* (chromosome 9) significant for V tortuosity, as well as *EFNB1*, the only gene found on the X chromosome, significant for V temporal angle. For males, three of the uniquely associated genes, *FMO3*, *FMO6P*, and *RBBP5*, clustered on the longer arm of chromosome 1 (1q); another three, *LAYN*, *SIK2*, and *PPP2R1B*, on chromosome 11l; two, *RAET1E* and *RAET1G*, were found, in close proximity, on chromosome 6, and one, *CIDP,* on chromosome 18. The traits for which these genes were significantly associated were A std diameter (genes on chr 1), A tortuosity (*LAYN*, *SIK2*), A/V ratio tortuosity (*PPP2R1B*), V central retinal eq (*RAET1E*, *REAT1G*), and V median diameter (*CIDP*). All gene results can be found here: [Significant Gene](https://docs.google.com/spreadsheets/d/1Uwjdnt4o-MKvGK1q7inMt56ng9mrgGGRz5qxMGlwbi8/edit?gid=1599558990#gid=1599558990).

### SNP-wise Heritability

We used heritability analysis to assess the genetic, SNP-wise, contribution to the phenotypic variance (*h*^2^_SNP_) for each IDP using GCTA’s GREML method (see Methods). We partitioned genetic variance and constructed separate genetic relationship matrices (GRMs) for each chromosome. *h*^2^_SNP_ was then estimated by jointly fitting each GRM with the multi-component GREML (MGRM) model. Individual chromosome *h*^2^_SNP_ results can further be found in **Suppl. Fig. 7a**.

Heritability estimates across all cohorts were largely consistent (**Fig. 4**), with mean *h*^2^_SNP_ across all 17 IDPs being very similar between the female, male, and combined cohorts 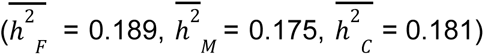. Albeit non-significant, we found *h*^2^_SNP_ to be slightly higher in females than in males for almost all traits, with the exception of V central retinal eq. Heritability for the X chromosome was negligible, with all *h*^2^_SNP_ < 1% across all IDPs and cohorts (**Suppl. Fig. 7b**). For detailed trait-wise heritability results, see **Suppl. Info. 4.**

**Figure 4.**
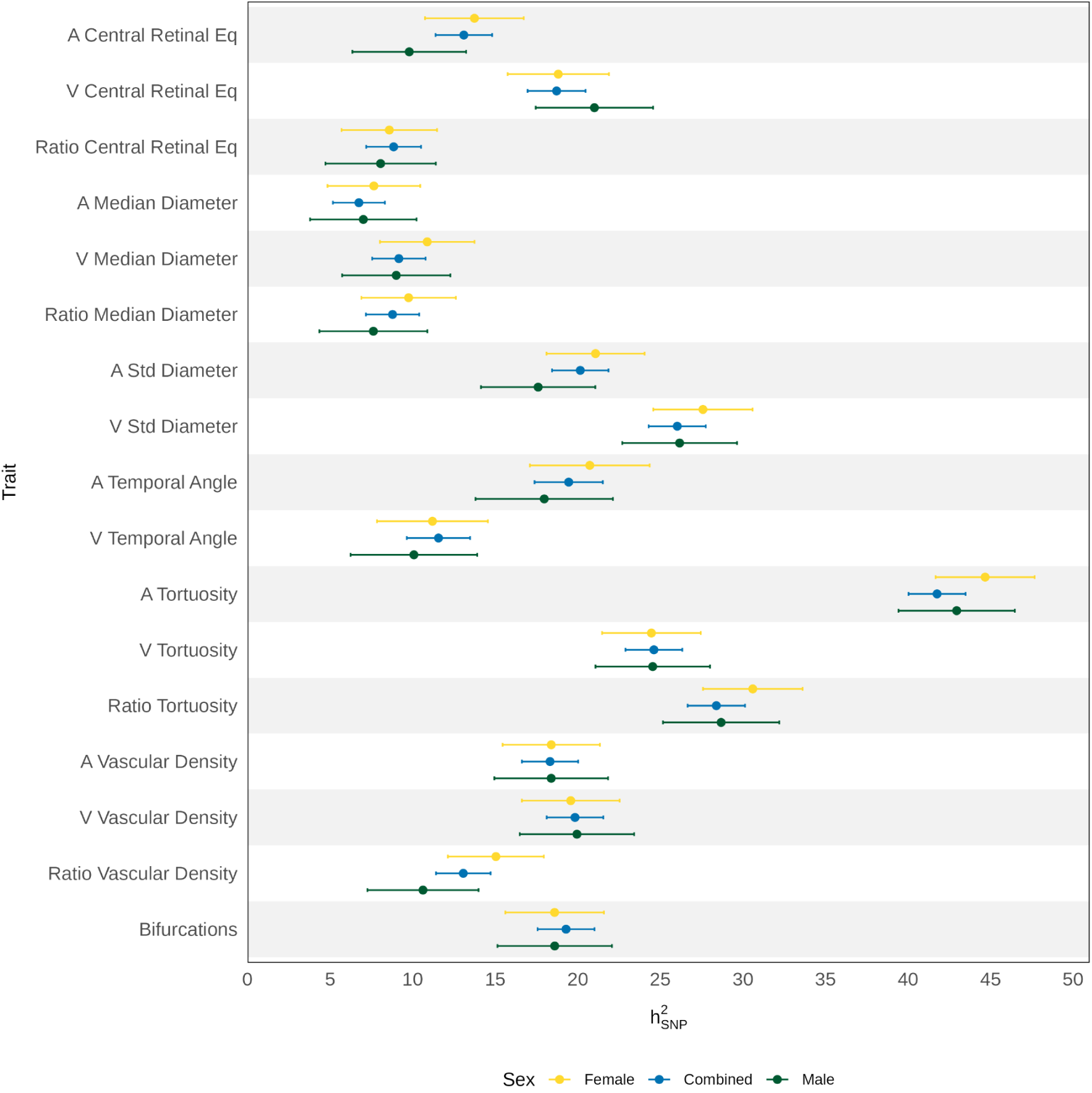
Comparison of h^2^_SNP_ estimates (in percent) for the 17 IDPs for each individual cohort. Error bars represent 95% confidence interval (CI) around the point estimate.

## Discussion

Sex has long been recognized as a critical variable in biomedical research, given its role in disease prevalence, expression, and progression for major causes of mortality such as cardiovascular and neurodegenerative diseases. Nevertheless, it remains notably underemphasized in the broader scientific literature. The same applies to the study of genetic architectures, where recent publications have shown modest, but significant sex-specific deviations in significant markers or heritability (Rawlik, Canela-Xandri and Tenesa, 2016; Bernabeu *et al*., 2021; Zhu *et al*., 2023). Here we expand on our previous work on retinal vascular phenotypes (Ortin Vela et al. (2024) focusing on sex-specific analyses. Specifically, we used interaction and stratified analyses for both disease and genome-wide regressions to assess significantly differing levels of association in the whole population but also to investigate sex-specific, independently adjusted effect sizes, and their divergence. We further included the X chromosome, often neglected in genetic analyses due to the analytical challenges it presents (Sun *et al*., 2023a), and are the first, to our knowledge, to report genetic associations on the X chromosome for vascular phenotypes.

While investigating differences in associations between IDPs, risk factors, and diseases, we found that the results presented an heterogeneous pattern across all traits or outcomes. Rather, we observed substantial sex differences in association of risk factors such as systolic blood pressure, hypertension, triglycerides, pulse rate, diabetes, and smoking, especially for bifurcations, A vascular density, and V vascular density. These differential results might indicate sex-differential impacts of these risk factors on microvascular phenotypes or reflect sex-specific lifestyle differences. This was partially expected: while the IDP distributions were found to be very similar between males and females, the distributions of risk factors notably diverged between males and females in some cases (**Suppl. Fig. 1b**). Similar to our findings in the risk factor and disease analysis, our survival analysis revealed substantial sex differences in the prognostic associations between IDPs and clinical outcomes, underscoring the importance of considering SABV in vascular research. In several cases, traits that appeared prognostic in the overall cohort were primarily driven by associations in one sex (e.g., V median diameter and A temporal angle for males; V vascular density and V tortuosity for females), while in others, stratified models uncovered associations that were absent in the pooled analysis (e.g., A/V ratio vascular density for death and A temporal angle for CVD death in males; A central retinal eq for CVD and A/V ratio tortuosity for death in females). These findings suggest that certain IDPs may have sex-specific prognostic value and highlight the potential for misinterpretation when sex differences are not explicitly accounted for. The variation we observed may reflect underlying biological differences in the microvasculature structure and function and indicate that IDPs may capture sex-specific pathways of vascular aging or disease progression. Besides supporting the biological plausibility of sex-specific microvascular phenotypes, this might also have practical implications for risk-stratification strategies and precision medicine.

We used the same two-faceted approach, interaction and stratification, to study sex differences in genetic architectures, demonstrating distinct but, to a certain degree, coherent signatures. Females seem to have more genetic signals associated with the IDPs, as evidenced by a trend towards higher heritability and larger numbers of SNP- and gene-associations. Interestingly, they also shared almost double the number of associated genes with the combined cohort as compared to males, suggesting that much of the genetic signal may be driven by females, or that it is more diluted in males due to environmental effects. This dilution may, in part, be due to the observed difference in risk factors, showing a trend towards higher blood pressure, BMI, and alcohol consumption, potentially reflecting poorer lifestyle. While the interaction analyses showed only six traits with rather few significant associations, one IDP, A/V ratio vascular density, exhibited genome-wide SNP x sex interactions with a highly significant signal on chromosome 15. Our sex-stratified analyses confirmed this sex-specific association, revealing a highly significant signal exclusively in females. The sex difference was further supported by the gene enrichment result, showing that *HERC2*, the gene within which these SNPs are located, was not significantly associated with this phenotype in males. Interestingly, the A/V ratio vascular density exhibited the largest sex difference in distribution among all traits and adjusting for the genetic effect of rs12913832 attenuated the phenotypic disparity almost completely. This SNP, located within an intron of *HERC2*, regulates the transcription of *OCA2*, a gene involved in melanin synthesis, and has been extensively studied for its strong association with eye color (Eiberg *et al*., 2008; Sturm *et al*., 2008; Visser, Kayser and Palstra, 2012; Meyer *et al*., 2020). Furthermore, it has recently been reported to be significantly associated with retinal pigmentation in a large GWAS study (Rajesh *et al*., 2025). Interestingly, this study did not find an effect of sex on retinal pigmentation (see their Suppl. Tab. 3), while previous work had suggested that sex may explain discrepancies in eye color prediction based on the *HERC2* locus (Martinez-Cadenas *et al*., 2013), although this effect seems population-specific (Pietroni *et al*., 2014). This raises the possibility that a sex-specific *HERC2* eQTL may contribute to differential melanin production in the retina, potentially altering vessel segmentation performance during image processing - either by enhancing artery visibility or reducing vein segmentation ability in females. Such an indirect effect may also explain other associations of retinal vascular properties with this locus, yet further work is needed to confirm this or rule out a direct effect of pigmentation on vessel properties, such as protection to structural damage from light.

We leveraged two complementary strategies to study sex differences in disease and genetic associations: an interaction approach, modeling the interaction between sex and phenotype or SNP, and a stratified approach, estimating effect sizes independently for each sex. Notably, the stratified approach fits a separate set of parameters for each sex, effectively doubling the number of estimated covariate effects compared to the interaction model. While both analyses showed rather concordant results, we found several IDP-disease association estimates to be unusually high compared to the Z test statistics and some GWAS results to be inflated for the interaction analyses. We hypothesize that these inflated estimates could reflect instability in the interaction models, potentially driven by multicollinearity or unaccounted confounding. This suggests that caution is needed when interpreting isolated interaction effects without complementary statistical approaches.

Building on the sex-stratified GWAS, our gene-level enrichment analysis revealed subtler but biologically relevant differences in genetic architecture between sexes. As with the SNP-level findings, the number of genes significantly associated with our IDPs was considerably lower in the sex-stratified analyses compared to the combined cohort. Nevertheless, we identified 12 genes uniquely associated with an IDP in one sex but not in the combined cohort, potentially highlighting sex-specific genetic mechanisms of microvascular morphology. In males, we observed unique associations with genes *FMO3*, *FMO6P*, and *RBBP5* (A std diameter), *LAYN* and *SIK2* (A tortuosity) and *PPP2R1B* (A/V ratio tortuosity), *RAET1E* and *RAET1G* (V central retinal eq), and *CIDP* (V median diameter). Variants of these genes have previously been implicated in blood lipid regulation and metabolism, including metabolite levels (*FMO3*, *FMO6P*) (Feofanova *et al*., 2020, 2023), triglycerides concentrations (*RBBP5*, *LAYN*, *PPP2R1B*) (Richardson *et al*., 2020; Ripatti *et al*., 2020; Sinnott-Armstrong *et al*., 2021), and protein levels (*LAYN*, *RAET1E*), (Emilsson *et al*., 2018; Western *et al*., 2024) but also aortic measurements and cardiovascular disease risk (*PPP2R1B*, *CIDP*) (Aragam *et al*., 2022; Francis *et al*., 2022; Verma *et al*., 2024). *RAET1E* and *RAET1G* further exhibit unique structural features within the *RAET1* family, encoding proteins that play important roles in immune function and inflammation. *RAET1E* in particular has been linked to vascular remodelling and inflammation through its ubiquitous expression in aortic endothelial cells and macrophage-rich lesion sites, indicating this gene to be a modifier of atherosclerosis susceptibility (Rodríguez *et al*., 2013). The presence of these significant associations suggests that, in males, vascular morphology might be more strongly influenced by lipid metabolism (i.e., LDL or very-low-density lipoprotein (VLDL), a major part of which is triglycerides), as well as by heightened inflammatory or immune responses. This is in line with previous evidence that males tend to exhibit higher LDL and triglyceride levels from adolescence onward, in part due to androgen-driven regulation of lipid pathways (Ciurtin, Taneja and Jury, 2023; Conlon *et al*., 2023). These sex differences may underlie the observed heightened susceptibility to atherosclerosis and cardiovascular disease, particularly in the younger male population. In females, we observed unique gene associations for *RBMXL1*, *GARNL3* (V tortuosity), and *EFNB1* (V temporal angle). Variants mapped to the two genes associated with V tortuosity have also been linked to metabolites, specifically histidine (Feofanova *et al*., 2023) and homovanillic acid (Luykx *et al*., 2014) metabolism - whose pathways intersect with hormonal regulation, particularly estrogen (Thompson and Moss, 1994; Mori *et al*., 2014; Yoest, Cummings and Becker, 2014), and are involved in inflammatory responses via histamine and dopamine. These neuromodulators can influence vascular structural stability through mechanisms such as histamine-induced vasodilation and increased permeability, or dopamine-mediated fluctuations in blood pressure and vascular resistance, suggesting a hormone-sensitive pathway contributing to sex-specific patterns of microvascular morphology in females. Mutations within *EFNB1*, the only gene located on the X chromosome in any of our gene enrichment analyses, are causative for craniofrontonasal syndrome (van den Elzen *et al*., 2014), leading to ophthalmic abnormalities, and have recently been implicated in retinal layer thickening (Jackson *et al*., 2025).

In contrast to the study by Ortin-Vela et al. (2024), which applied LD score regression (Bulik-Sullivan *et al*., 2015) for heritability estimation, we used GCTA’s Genomic-relatedness-based Restricted Maximum-Likelihood (GREML) methodology (Yang *et al*., 2010, 2011). This method allowed us to model the phenotypic similarity between unrelated individuals as a function of their genetic relatedness and partition the phenotypic variance into genetic and residual components for the autosomes and the X chromosome without relying on GWAS summary statistics or an LD reference panel. In our analysis, *h*^2^_SNP_ estimates in the combined sample are comparable to or higher than in the previous study, adding up to ∼17% more explained variance for A tortuosity (for a comparison see **Suppl. Fig. 7d**). Considering that the reference panel used in Ortin-Vela et al. (2024) was the HapMap 3 1000 Genomes project (EUR), heritability might have been underestimated by LDSC, a pattern which has been noted in a previous comparison (Ni *et al*., 2018; Srivastava, Williams and Zhang, 2023). By stratifying our sample, we further observed higher *h*^2^_SNP_ for the microvasculature in females, with absolute values generally more similar to those of the combined cohort, suggesting that genetic variance in this sex might contribute more substantially to the proportion of explained phenotypic variance. For A tortuosity and the A/V tortuosity ratio, we found higher *h*^2^_SNP_ estimates for both sexes. This pattern – sex-stratified estimates exceeding those of the combined sample – may reflect sex-specific genetic effects that are masked when analyzed jointly or reduced environmental heterogeneity within each sex, increasing the proportion of phenotypic variance attributed to genetics. Furthermore, our results of >40% for A tortuosity in all cohorts exceed previously reported heritabilities with higher estimates observed only in one other population-based study (Jiang *et al*., 2023) as well as in twin (82%) (Taarnhøj *et al*., 2008) or small family-based (55%) (Kirin *et al*., 2017) studies.

Whereas this study implemented multiple new approaches, i.e., sex-specific analyses, Cox modeling correcting for covariates used in the Framingham model, up-to-date GWAS methodology, heritability estimation directly from genotypes, and X chromosome inclusion, we would like to point out several important limitations: first, our cohort was a subset of the UKB and thus encompasses predominantly individuals of white British ancestry of middle to old age. Large volunteer-based cohorts like the UKB have been shown to exhibit selective participation, with respondents, on average, being healthier and of higher socio-economic status than the overall population, potentially creating bias in disease or phenotype-genotype associations (Fry *et al*., 2017; Schoeler *et al*., 2023; van Alten *et al*., 2024). Second, we relied on the phenotypes extracted through the automated pipeline described in Ortin-Vela et al. (2024), which might be inherently biased because of the way the underlying deep-learning segmentation was trained. Depending on the original dataset used to train the algorithm, performance might be poor in specific subgroups, i.e., different sexes, ages, or ancestries, an issue known as “unfairness” in deep-learning-based image analysis (DeCamp and Lindvall, 2020; Liu *et al*., 2023). Third, we note that while we tried to mitigate problems associated with self-reported measures by primarily using electronic health record data, these sources may still exhibit systematic bias, particularly in the female population: extensive literature documents gender bias in CVD diagnosis and treatment, highlighting that women are frequently underdiagnosed due to “atypical” presentations, their symptoms being dismissed or misattributed, and a lower likelihood of receiving comprehensive diagnostic testing or preventative intervention (Bairey Merz, Andersen and Shufelt, 2015; Desai, Munshi and Munshi, 2021; Al Hamid *et al*., 2024). Lastly, we note that the different frequencies of cardiovascular events and deaths between the male and female cohorts may introduce additional sources of bias. Given that statistical power is tied to event rates, the lower number of events observed in females could limit the ability to detect associations, affecting the precision of point estimates, the performance of prognostic models, and the reliability of statistical significance testing.

In conclusion, this study provides evidence for sex-specific differences in diagnostic and prognostic effect estimates, as well as in genetic architecture of parameters describing the retinal microvasculature. While trait distributions differed only minimally between females and males, both interaction and sex-stratified analyses revealed clear sex-specific differences in how traits relate to cardiometabolic risk factors and diseases. Sex-stratified survival analysis for major cardiovascular mortality drivers as well as death outcomes further uncovered a distinct pattern with some traits showing significant prognostic value for mortality risk evaluation in one sex, potentially driving the signal in the overall population. Unlike traditional CVD risk markers, such as blood lipid or pressure measurements, which might fluctuate substantially over short periods of time, microvascular traits likely provide a more stable reflection of vascular health. Despite being influenced by lifestyle factors, processes of microvascular remodeling take place over time and therefore capture cumulative stress but also, potentially, differences in genetic susceptibility. Our sex-stratified genetic analyses underline this point, showing subtle differences in genetic architecture, heritability, and significant gene results, indicating sex-specific mechanistic pathways or environmental exposures contributing to microvascular morphology. These findings would have remained undetected in sex-agnostic analyses, highlighting the necessity of incorporating sex-specific approaches to fully understand complex trait biology and improve precision in cardiovascular risk prediction.

## Methods

### Phenotypic data preparation

The data used here originates from the study of Ortin-Vela et al. (2024) and consists of 17 image-derived phenotypes (IDPs) from the retinal vasculature for a subset (n=71,494) of the UK Biobank (UKB). More detailed descriptions of the IDPs, including their derivation, are provided in Ortin Vela et al. (2024). The UKB is a large-scale biomedical database and research resource containing genetic, lifestyle, and health information from half a million UK participants. In addition to the quality control employed in Ortin-Vela et al. (2024), individuals with sex chromosome aneuploidies (UKB 22019) as well as those whose genetic sex (UKB 22001) did not match their reported sex (UKB 31) were excluded, resulting in a total of 68,623 individuals (Female = 37,080, Male = 31,543). Sex chromosome aneuploidies can lead to atypical phenotypic presentations, the implications of which on the microvasculature remain poorly understood. Additionally, since discrepancies between genetic and reported sex can introduce confounding effects related to the complex interplay between biological sex and gender, further complicating trait-disease associations, we excluded all such cases.

### Disease Association

Each retinal IDP was tested for association with a number of diseases, which can generally be divided into eye and cardiovascular diseases (CVD), and systemic risk factors known to affect disease susceptibility. Individual-level disease and risk factor information was collected from the UKB and the corresponding fields as well as further information can be found in **Suppl. Info 3**. Information provided through health-related records (Category 100091), biological samples (Category 100078), Physical (100006) and Eye measures (Category 100013) were prioritized over those obtained through the assessment center touchscreen Questionnaire (Category 100025) or Verbal Interview (Category 100071) as self-reported measures are often unreliable due to high reporting errors or inconsistencies (Modin *et al*., 2017; Schoeler, Pingault and Kutalik, 2024). Exceptions were made for alcohol intake frequency, drinking (i.e., never, previous, current), and smoking status as well as pack-year smoking. As studies have shown the prevalence of undiagnosed diabetic or pre-diabetic conditions and associated health complications (Dankner *et al*., 2009; Pierce *et al*., 2009), diabetes status was determined using a combination of variables: an official diagnosis based on the date E10 first reported (insulin-dependent diabetes mellitus), HbA1C readings of larger or equal to 48 mmol/mol, or glucose levels of larger or equal to 11.1 mmol/l (Qiao *et al*., 2023; Lugner *et al*., 2024).

Sex-stratified multiple linear regression models were employed for the continuous risk factors and multiple logistic regression models for the disease traits, including in all cases a number of covariates known or hypothesized to confound retinal vascular variability: age, assessment center, image instance, spherical and cylindrical power, genotype measurement batch as well as the first 20 genomic PCs (for more information, see **Suppl. Table 3.1**). Prior to model fitting, all IDPs were normalized (z-scored) individually for the female and male cohorts, and all diseases collected as “date first diagnosed” were binarized. Standardized beta coefficients and odds ratios were extracted from the regression models, and significance was determined using two Bonferroni thresholds, i.e., p < 0.05/N_Tests_ (denoted as *) or p < 0.001/N_Tests_ (denoted as **), where N_Tests_= N_IDPs_ × N_Diseases_ (**Suppl. Fig. 3**).

To assess differences in risk and disease association between females and males, Z-tests (or Clogg tests) (Clogg, Petkova and Haritou, 1995) were used. Z-tests compare two beta coefficients or odds ratios from different models (or the same model under different conditions) to determine whether their difference is significant relative to their standard error: 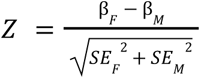. A positive Z score will correspond to a higher association in females, while a negative Z score will signal a higher association in males. Significance was assumed for p < 0.05 (*) with multiple testing thresholds set to p < 0.05/ N_IDP_ (**), and p < 0.05/ N_IDPs_ x N_Diseases_ (***).

### Survival Analysis

Six distinct endpoints were defined for the survival analysis: all-cause mortality (death), cardiovascular mortality (CVD death), cardiovascular events (CVD), Ischaemic Heart Disease (IHD), Myocardial Infarction (MI), and stroke. Cardiovascular-related death was defined using the reported ICD-10 codes. Time to event (TTE) was defined as the days between the date of enrollment in the study and the date of the reported event. If the subject experienced multiple events, the first event after the imaging visit was used. The risk factors included in the model matched those in the Framingham score (Wilson *et al*., 1998), i.e., age, SBP, total as well as HDL cholesterol, BMI, smoking status, diabetes diagnosis, and hypertension treatment (i.e., medication). Hypertension treatment was defined by aggregating male- and female-specific medication and filtering for “blood pressure medication”. Diagnosis of previous cardiovascular events was further included due to an increased risk of recurrence (Lassenius *et al*., 2021). All the fields used for the analysis and their descriptions can be found in **Suppl. Tables 3.2–3.4**.

### Genome-wide association study (GWAS)

GWAS was performed using REGENIE, which implements a robust two-step whole-genome regression approach accounting for underlying population structure (Mbatchou *et al*., 2021). Prior to the analyses, the genotypic data were prepared as recommended for the UKB, creating one file for all chromosomes, including only directly genotyped markers with further quality control, including MAF 0.01, MAC 100, genotype missingness 0.1, and HWE 1e-15. As there is no consensus on quality control for the X chromosome, due to its unique biological features, we applied several controls tailored to our study: the X chromosome was prepared independently in each sex, relaxing the genotype missingness for males to 0.2 and excluding all ampliconic regions (Gao *et al*., 2015; Khramtsova *et al*., 2023). For the X chromosome, the HWE filter was further removed as standard HWE tests are not directly applicable to hemizygous males and may lead to the unnecessary exclusion of variants (Waples, 2015; Graffelman and Weir, 2016).

The raw phenotypic data were transformed using rank-based inverse normal transformation. The covariates included in the GWAS were age at assessment center visit, age at assessment center visit squared, assessment center location, instance, spherical power, spherical power squared, cylindrical power, cylindrical power squared, genotype measurement batch, and the first 20 genomic PCs.

Association testing was performed for all three cohorts (female, male, combined) with per-trait sample sizes varying due to different missingness patterns. The sex-stratified cohorts’ sample sizes ranged from 25,069 or 29,790 (A temporal angle) to 31,332 or 36,802 (A and V vascular density) individuals for males and females, respectively. The Manhattan and QQ plots for all individual traits for each cohort can be found in **Suppl. Fig. 3 a-f**.

### Meta Analysis

The sex-stratified results for each retinal vascular trait were meta-analyzed using the open-source software GWAMA (Mägi and Morris, 2010), which implements a fixed effect model combining all allelic effects for each individual SNP, weighted by their inverse variance, as well as tests of between-study heterogeneity. The *I*^2^_j_ statistic, which quantifies the variation in SNP effects across all included studies over that expected by chance (Higgins and Thompson, 2002; Huedo-Medina *et al*., 2006), was used to assess SNP-wise heterogeneity between Female and Male results, with a general cutoff of 75% being considered as high variability. Pooled SNP effect sizes, directions, and heterogeneity estimates were extracted from the resulting summary statistic. *I*^2^ results were interpreted in the context of individual cohorts and pooled effect size estimates, considering limited power when only considering two studies. Individual IDP meta-analysis results (Manhattan and QQ) can be found in **Suppl. Fig. 3 g-h**.

### Gene Scoring

SNP effect sizes of all GWAS summary statistics were aggregated for gene scoring using PascalX (Krefl, Brandulas Cammarata and Bergmann, 2023). PascalX employs the sum of χ^2^ method to test for GWAS gene enrichment, an algorithm originally introduced by Lamparter et al. (2016). The software was kindly modified by D. Krefl to allow for X-chromosomal gene scoring. The UK10K (UK10K Consortium *et al*., 2015) panel was used for gene scoring on the autosomes while separate X chromosome panels were constructed for females and males, each using 5,000 unrelated white individuals of the respective sex from the UKB. Protein-coding genes as well as lincRNAs available at Ensembl’s Biomart were scored (n = 19936) using the approximate “saddle” method and default options for the gene window (50 kb), MAF (0.05), and VARCUTOFF (0.99). Cross GWAS coherence enrichment tests (Krefl and Bergmann, 2022) were employed for sex-specific summary statistics to reveal differences in within-trait gene effect direction (coherent vs anti-coherent). Anti-coherence results, while not significant, can be found in **Suppl. Fig. 8**. In all analyses, genes were considered significant after multiple testing corrections (p = 0.05/ N_genes_).

### Heritability

SNP-based heritabilities (h^2^_SNP_) were estimated using the GCTA (Genome-wide complex trait analysis) software package for each cohort independently. In particular, the genomic-relatedness-based restricted maximum-likelihood (GREML) method (Yang *et al*., 2010, 2011) was used to estimate the proportion of variance of the IDP explained by the genetic data. The genetic relationship matrix (GRM) was constructed independently for every cohort per chromosome using only directly genotyped SNPs with an MAF > 0.01 to reduce computational burden and minimize bias due to imputation quality. For the X chromosome, equal genetic variance between Females and Males was assumed (Keur *et al*., 2022). The h^2^_SNP_ results were obtained using the multi-component GRM (MGRM) model, using one genetic relationship matrix for all autosomes and an additional one for the X chromosome. We further estimated heritability for each individual chromosome (**Suppl. Fig. 5**).

## Data Availability

Comprehensive disease diagnoses, risk factor data, and raw phenotypic data are available under restricted access from the UK Biobank and can be obtained upon project approval (https://www.ukbiobank.ac.uk/enable-your-research/apply-for-access). GWAS summary statistics generated in this study will be made publicly available via the NHGRI-EBI catalog of human genome-wide association studies (https://www.ebi.ac.uk/gwas/home) upon publication.

## Code Availability

All code will be made available on github upon publication.

## Acknowledgments

The authors would like to thank Daniel Krefl for his contribution in implementing X chromosomal gene scoring in PascalX.

*Funding*: Supported by the Swiss National Science Foundation, Sinergia grant no. CRSII5 209510, “Connecting properties of the micro- and macrovasculature from multimodal imaging through genetics and deep learning to better understand vascular pathomechanisms and predict disease risk”.

## Author contributions

LB conceived and designed the study, performed disease association and genetic analyses, and drafted the manuscript. DB contributed to disease association, performed survival analyses, and assisted in manuscript drafting. SvB provided supervision, conceptual guidance, and critical manuscript revision. OT offered scientific advice on phenotypic analyses and contributed to manuscript revision. MB provided scientific advice on genetic analyses, reviewed the manuscript, and supplied code. DP, IQ, SaB, and II reviewed and provided comments on the manuscript.

