## Supplementary Figures for "Sex-specific disease association and genetic architecture of retinal vascular traits"

**Supplementary Figures 1**: Phenotypic trait and disease distributions between the sexes.

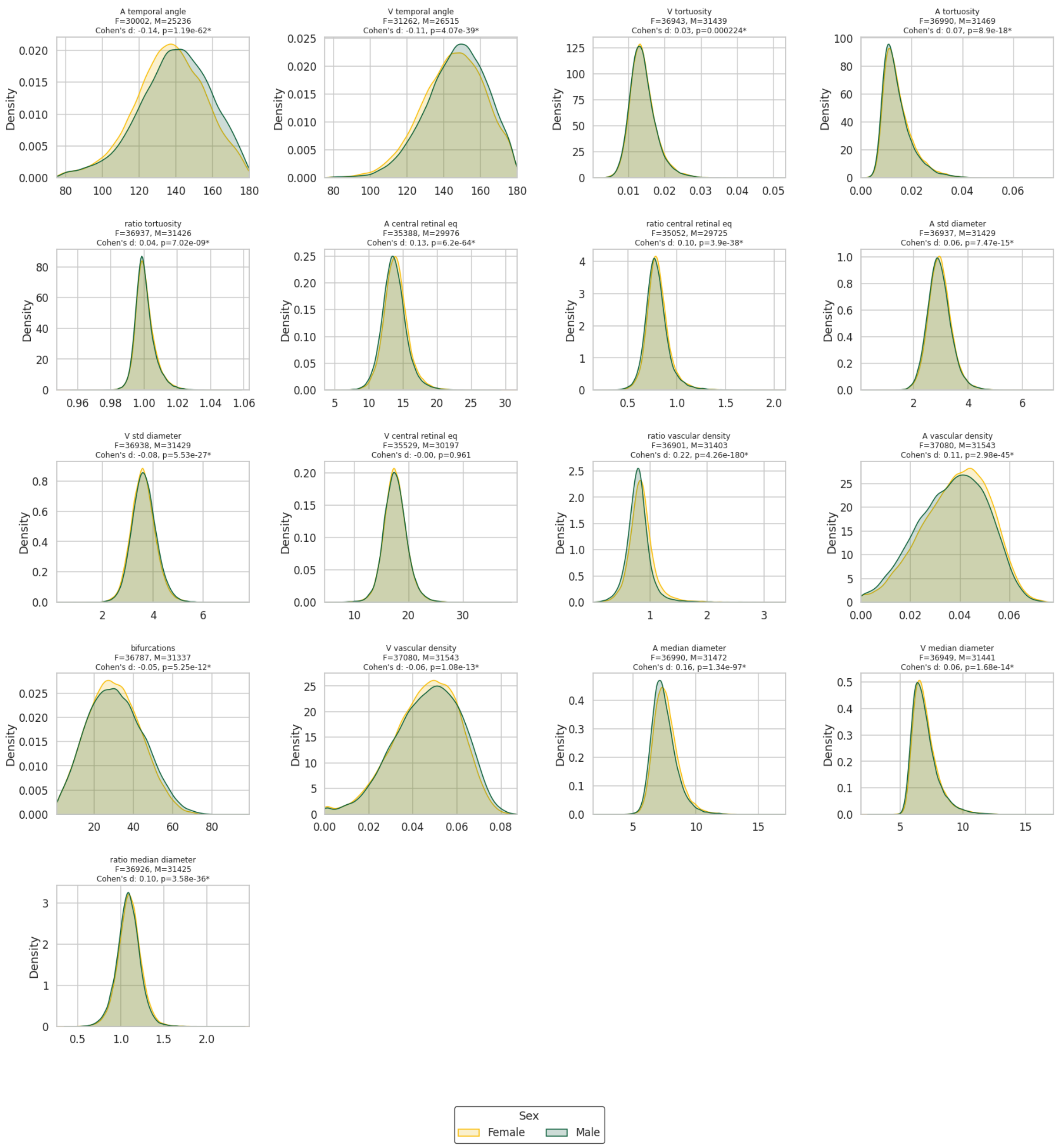


a) Distributions for all 17 retinal phenotypes. Cohen's d and associated p values reported in each subplots header with statistical significance indicated as "∗" after Bonferroni correction (∗ = $p < 0.05/N_{IDPs}$).

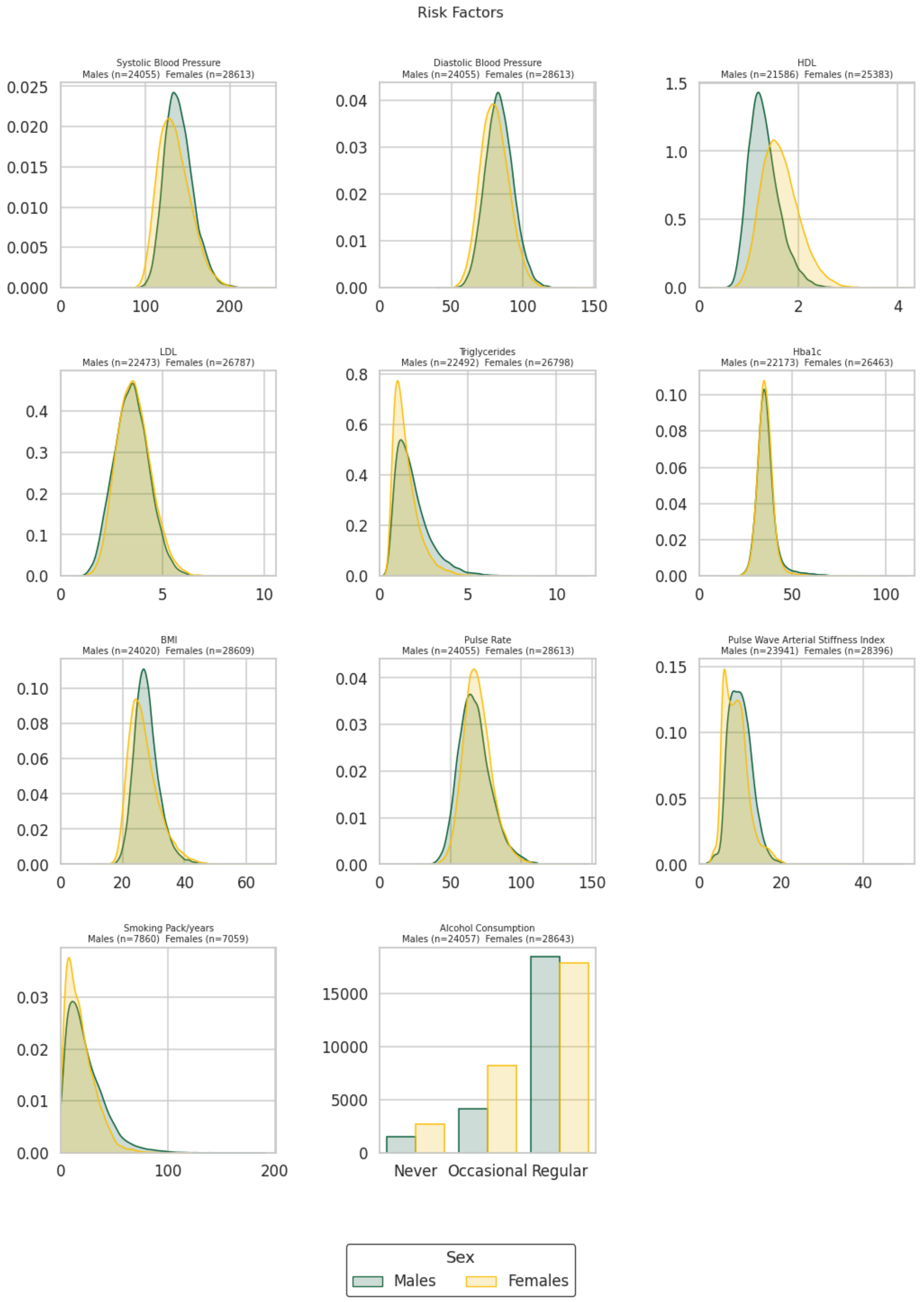


b) Distributions for all risk factors considered in the linear regression. Outlier values (10 std +/- mean) removed.

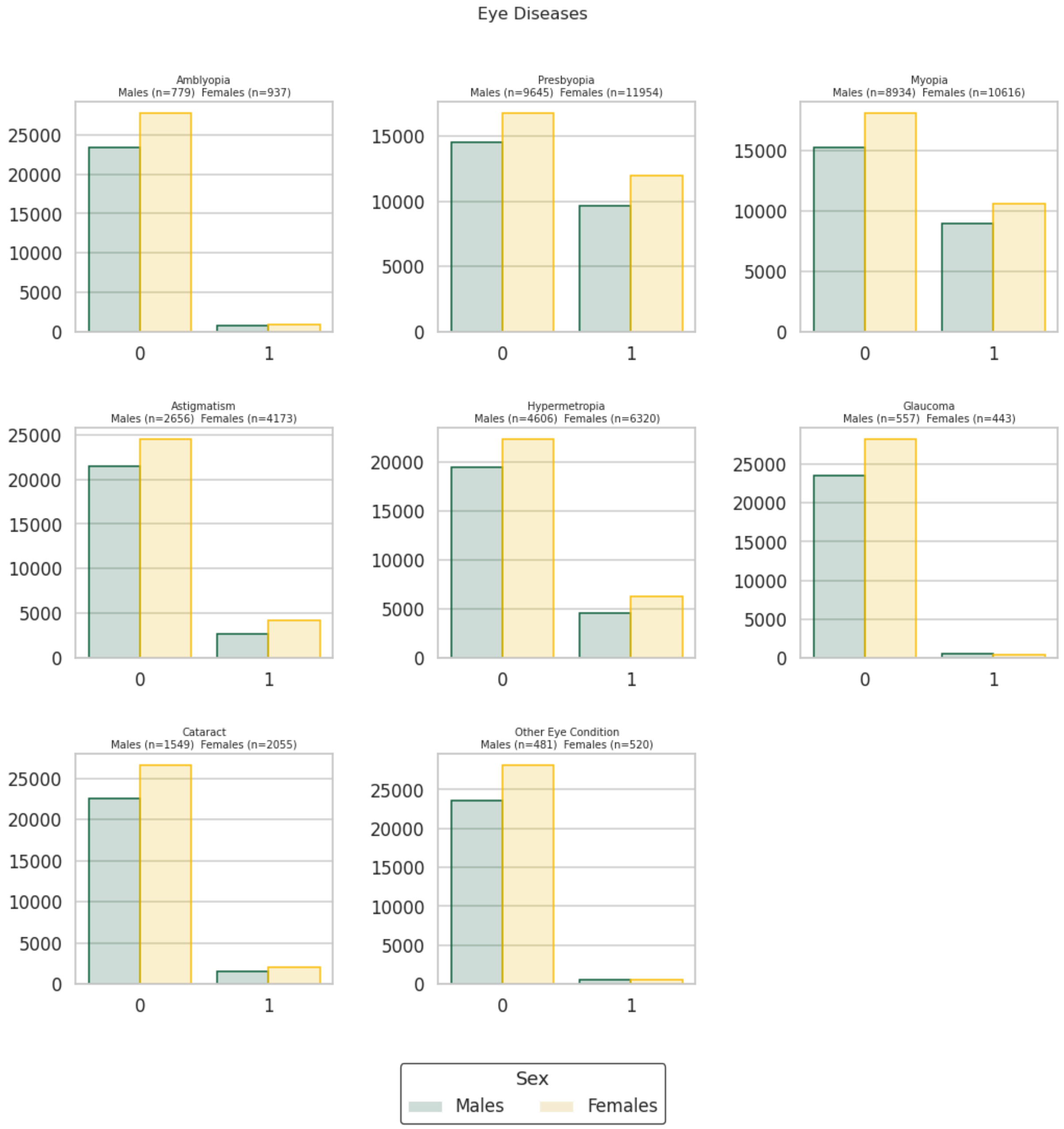


c) Distributions for all eye diseases considered in the logistic regression.

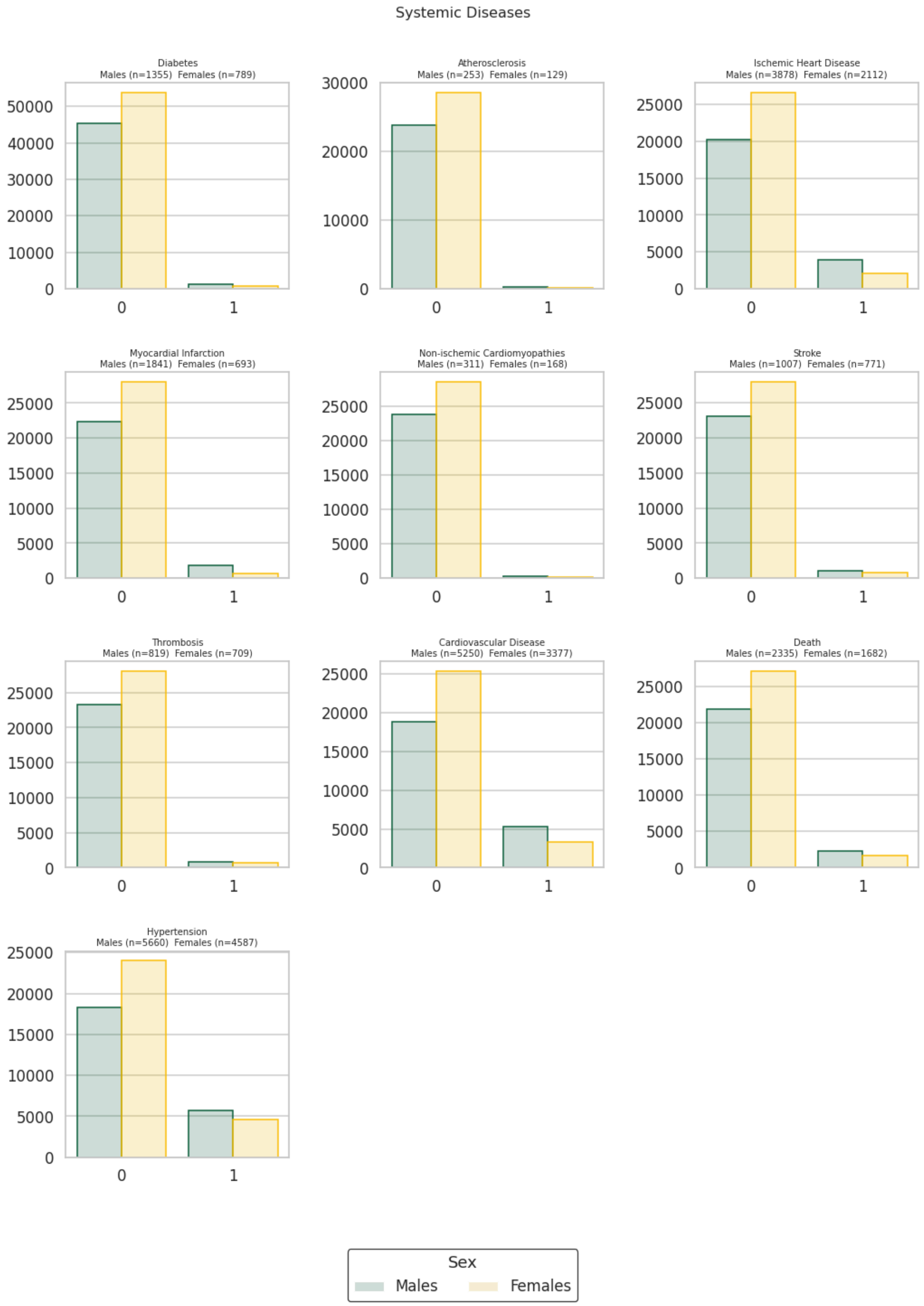


d) Distributions for all systemic diseases considered in the logistic regression.

**Supplementary Figures 3:** Sex-specific disease association analyses

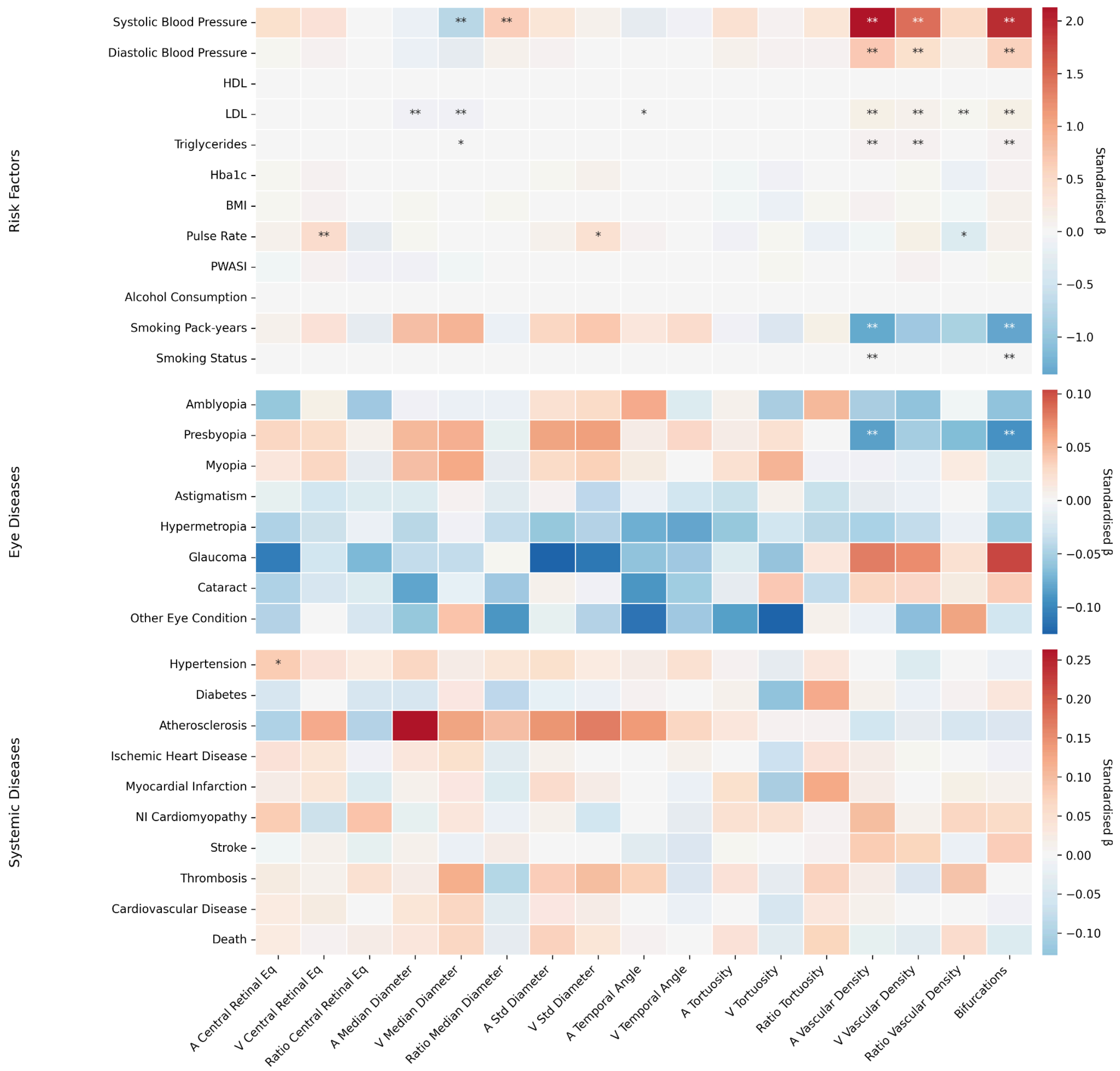


a) Trait - Disease association for IDPxSex interaction. Asterisks indicate the level of statistical significance after Bonferroni correction (* = $p < 0.05/N_{Tests}$, ** = $p < 0.001/N_{Tests}$, where $N_{Tests} = N_{IDPs} \times N_{Diseases}$ where $N_{Diseases}$ was considered the sum of all dependent variables used in the linear (n = 11) and logistic regressions (n = 18).

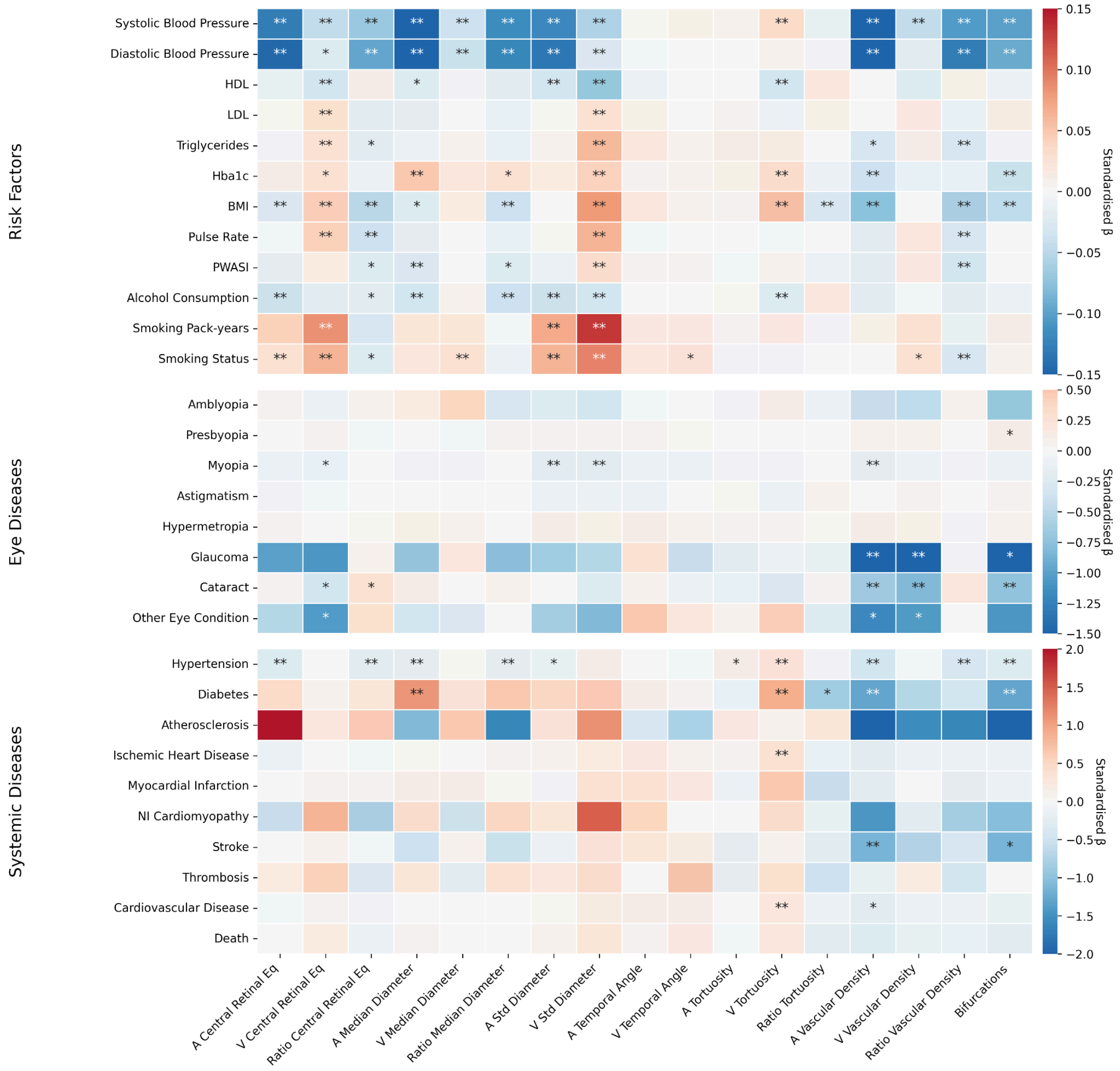


b) Trait - Disease association for females. Asterisks indicate the level of statistical significance after Bonferroni correction ($* = p < 0.05/N_{Tests}$, $** = p < 0.001/N_{Tests}$, where $N_{Tests} = N_{IDPs} \times N_{Diseases}$ where $N_{Diseases}$ was considered the sum of all dependent variables used in the linear (n = 11) and logistic regressions (n = 18).

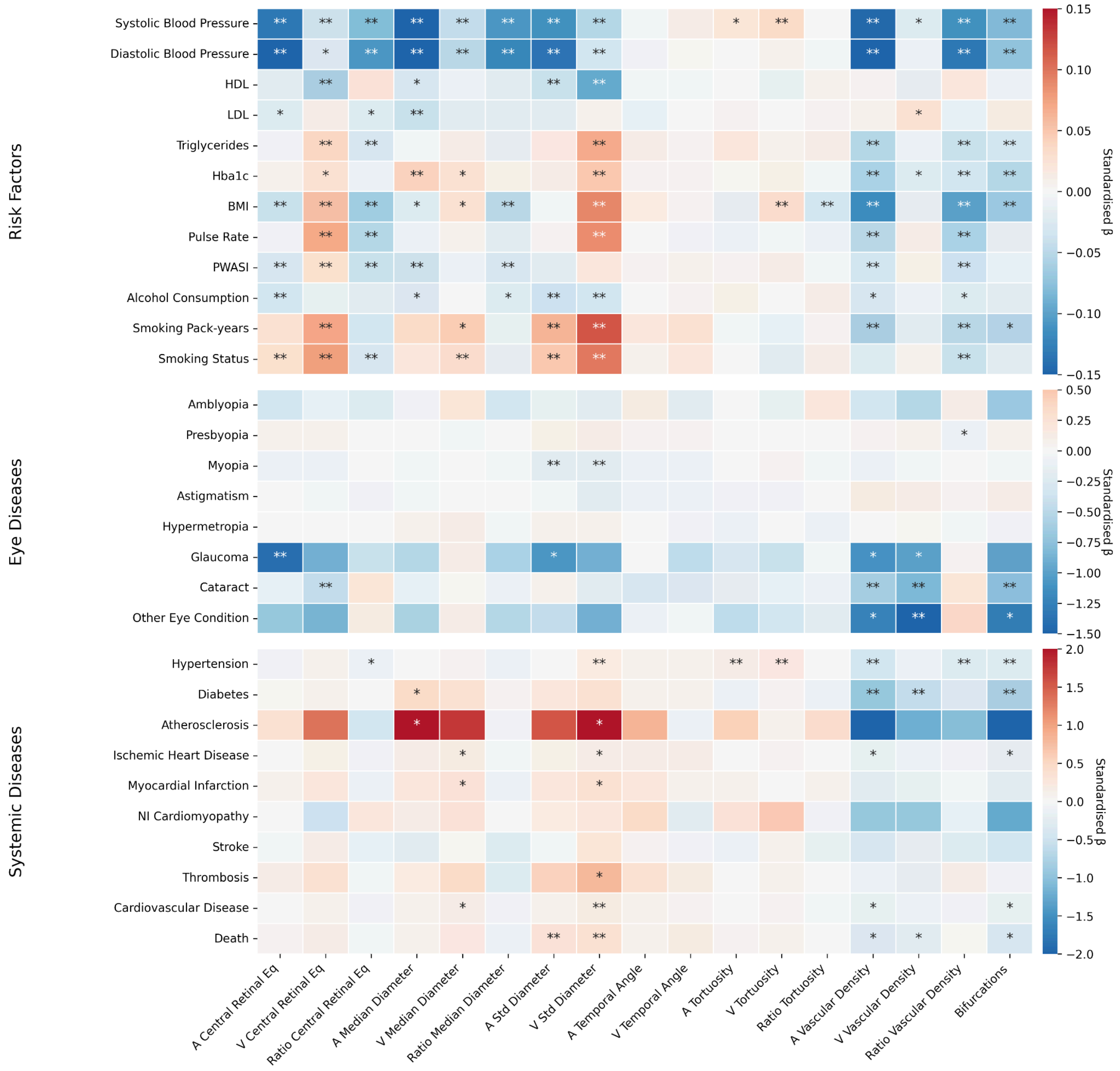


c) Trait - Disease association for males. Asterisks indicate the level of statistical significance after Bonferroni correction (∗ = $p < 0.05/N_{Tests}$, ∗∗ = $p < 0.001/N_{Tests}$, where $N_{Tests} = N_{IDPs} \times N_{Diseases}$ where $N_{Diseases}$ was considered the sum of all dependent variables used in the linear (n = 11) and logistic regressions (n = 18).

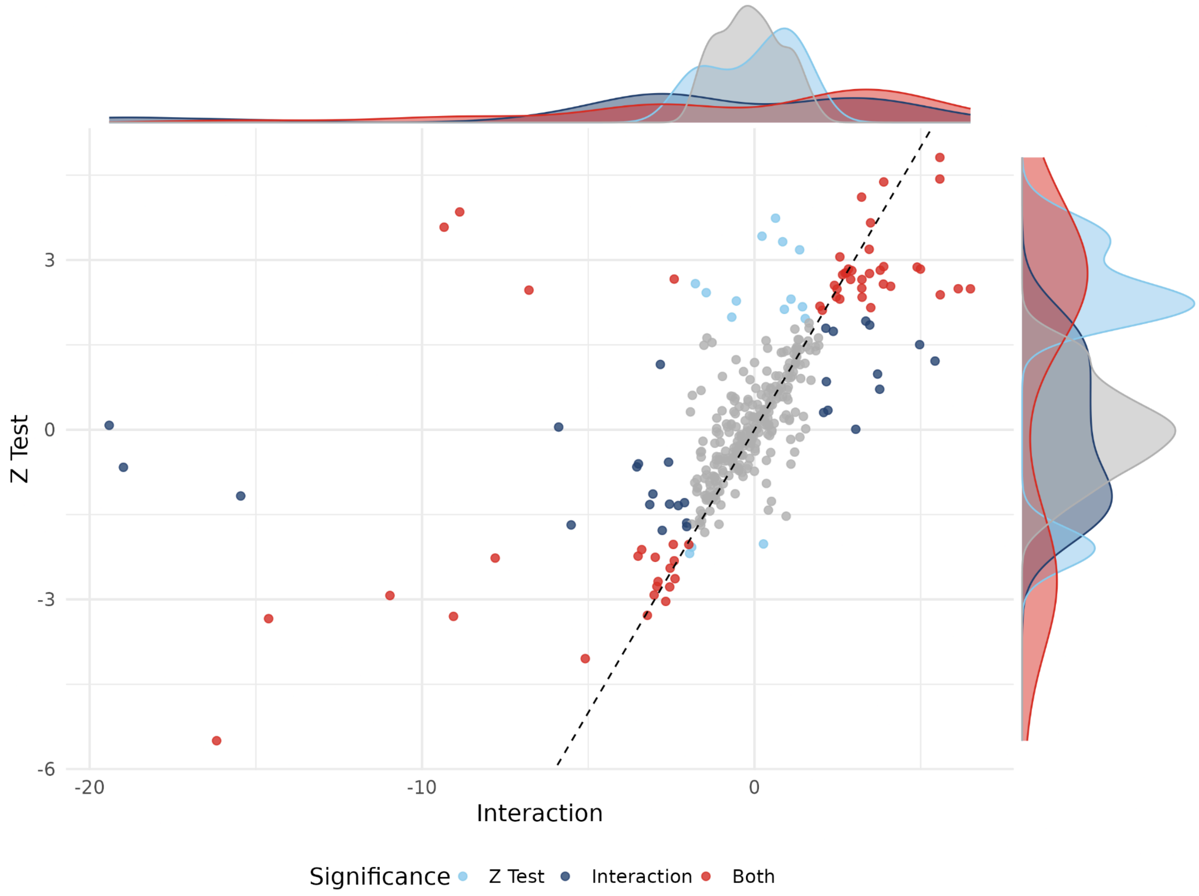


d) Comparison of interaction effects and Z-test results. Statistical significance considered at a threshold of 0.05. Interaction effect estimates flipped for concordant effect direction between analyses.

**Supplementary Figures 4:** Survival Analysis

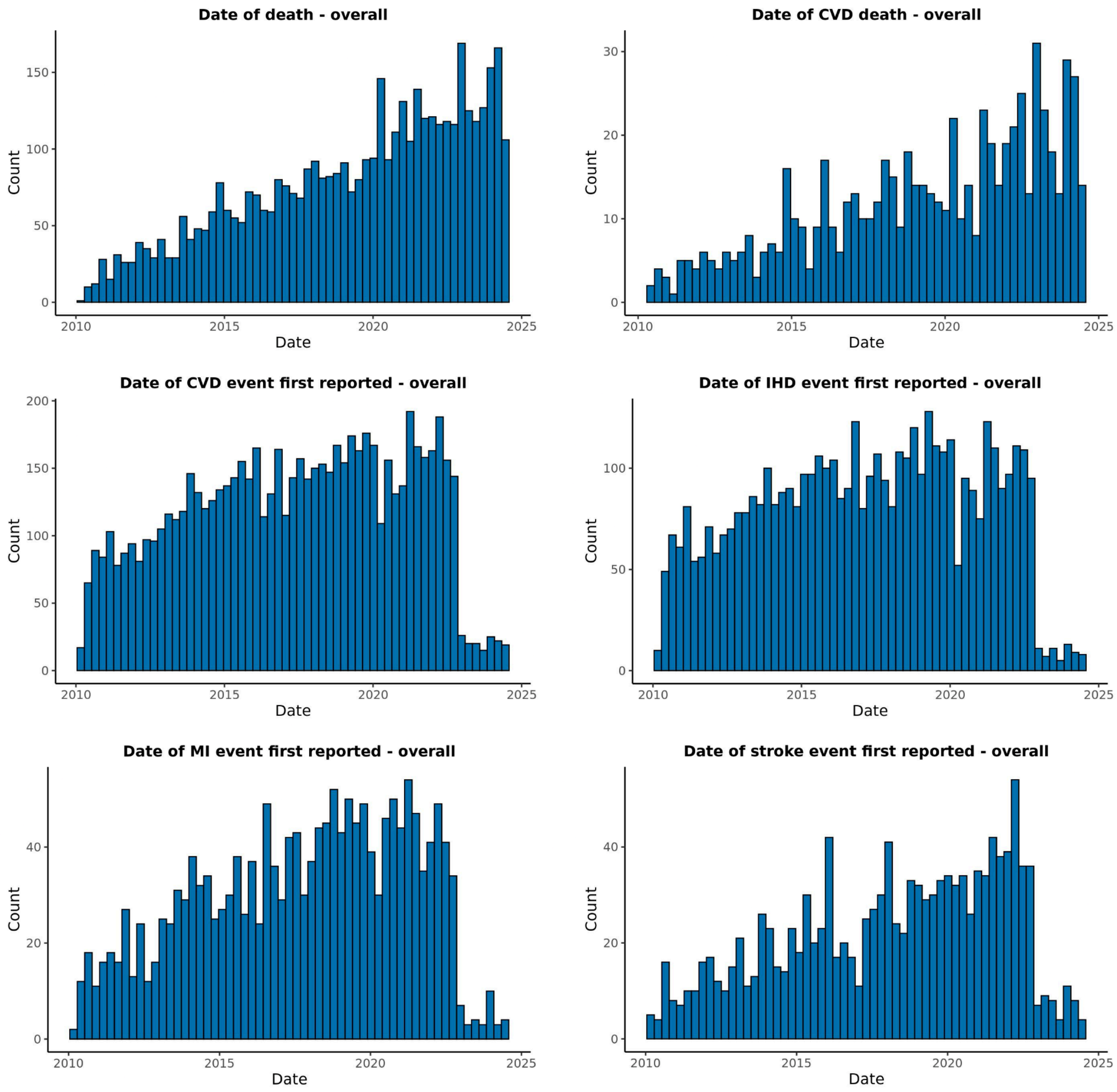


a) Distribution of date of event for the selected endpoints for the combined cohort. Only events after colour fundus image acquisition were considered. If one subject experienced multiple events, the first event after the imaging visit was used.

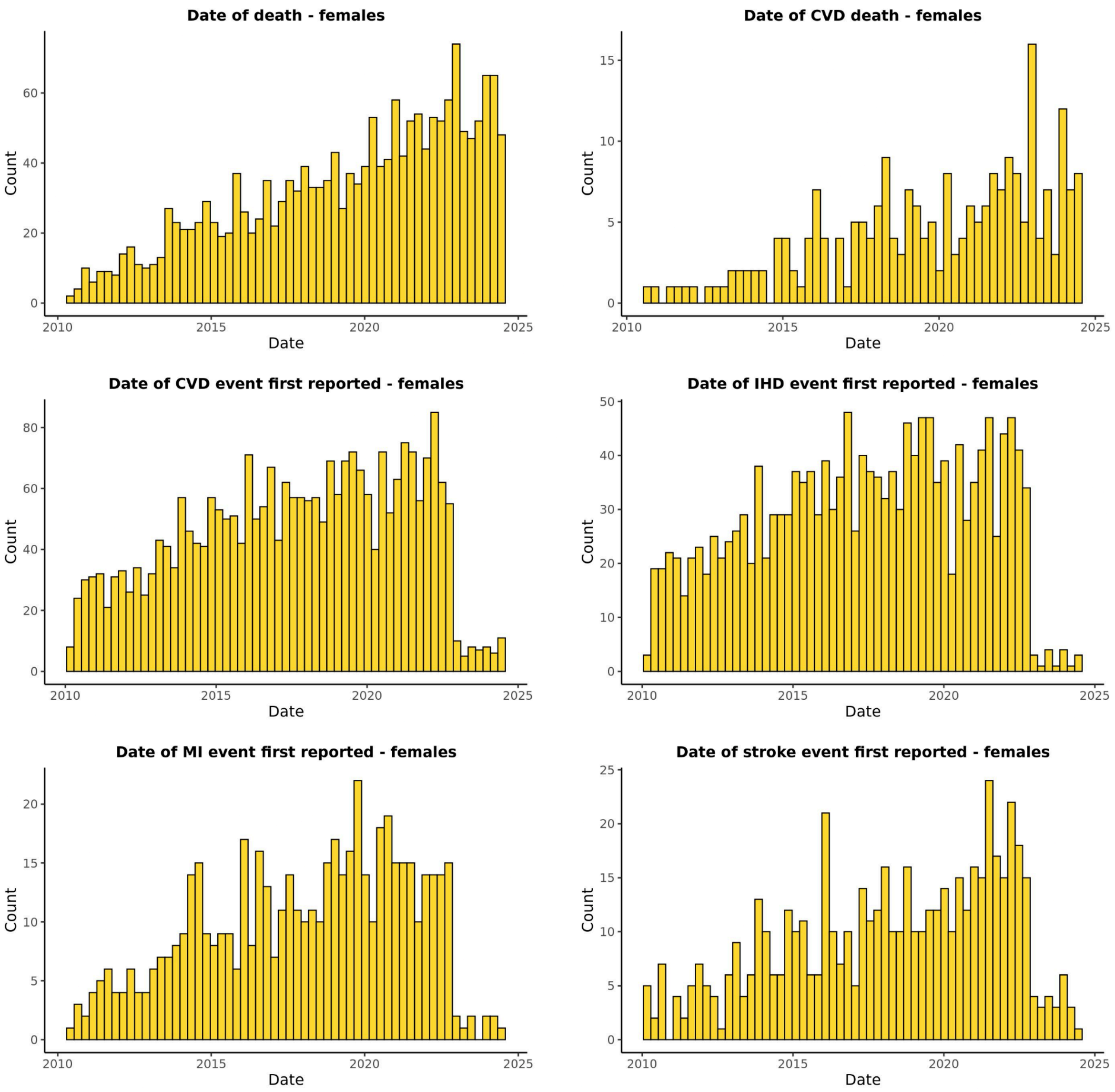


b) Distribution of events for the selected endpoints for the female cohort.

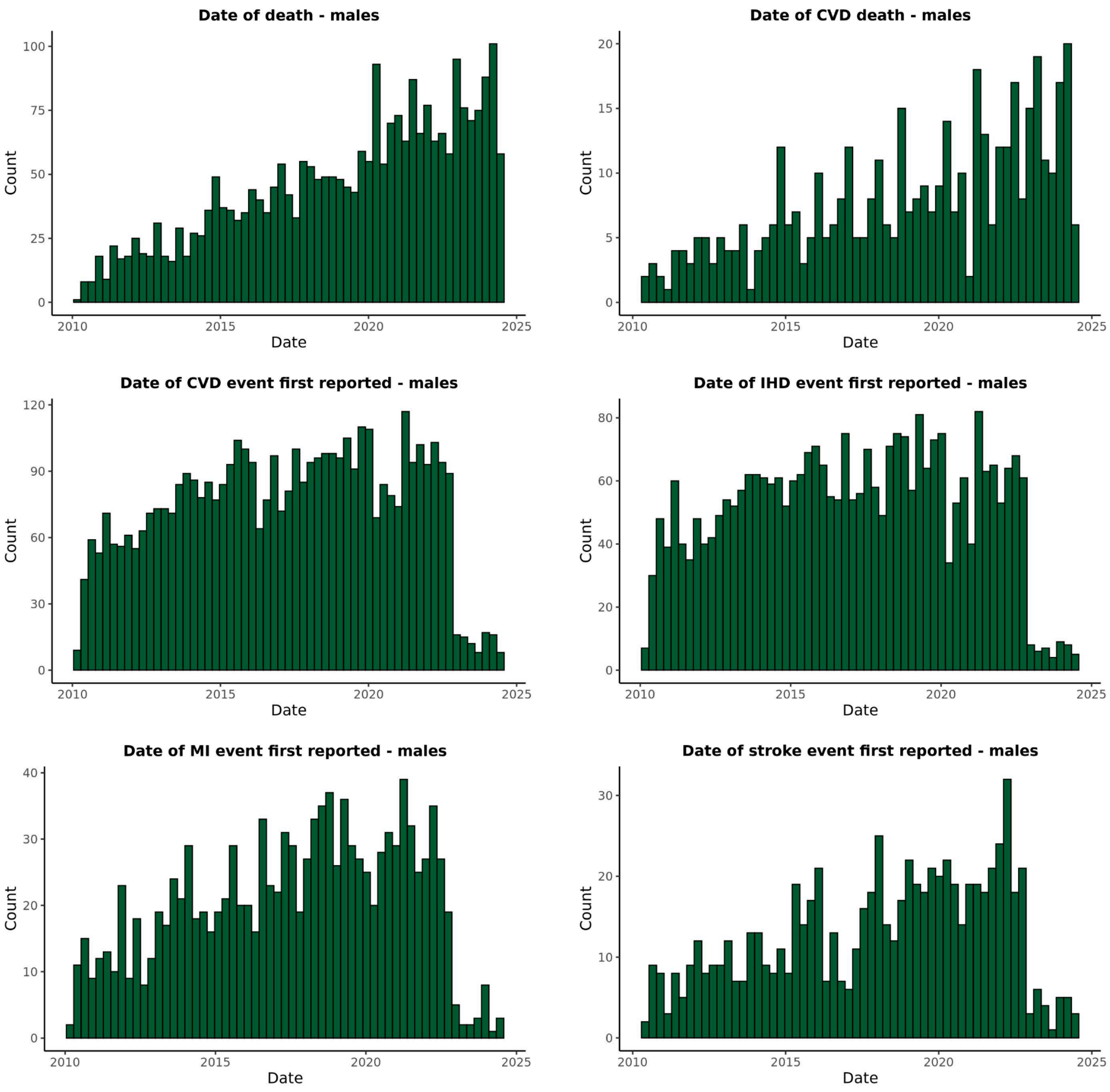


c) Distribution of events for the selected endpoints for the male cohort.

**Supplementary Figures 5**: Manhattan and QQ plots

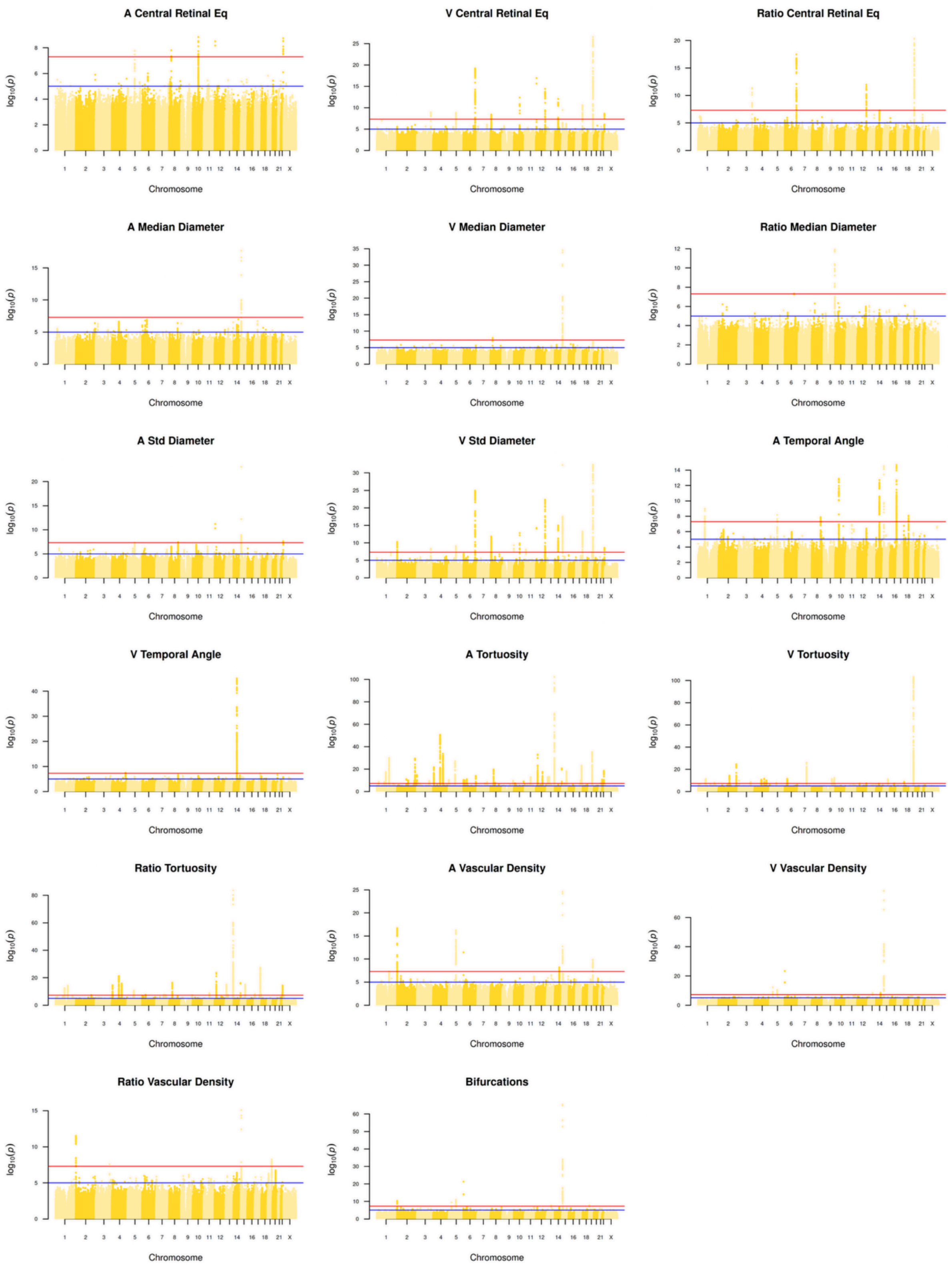


a) Manhattan plots of all 17 retinal traits for Females. Genome-wide significance (5e-8): red line, near genome-wide significance (1e-5): blue line.

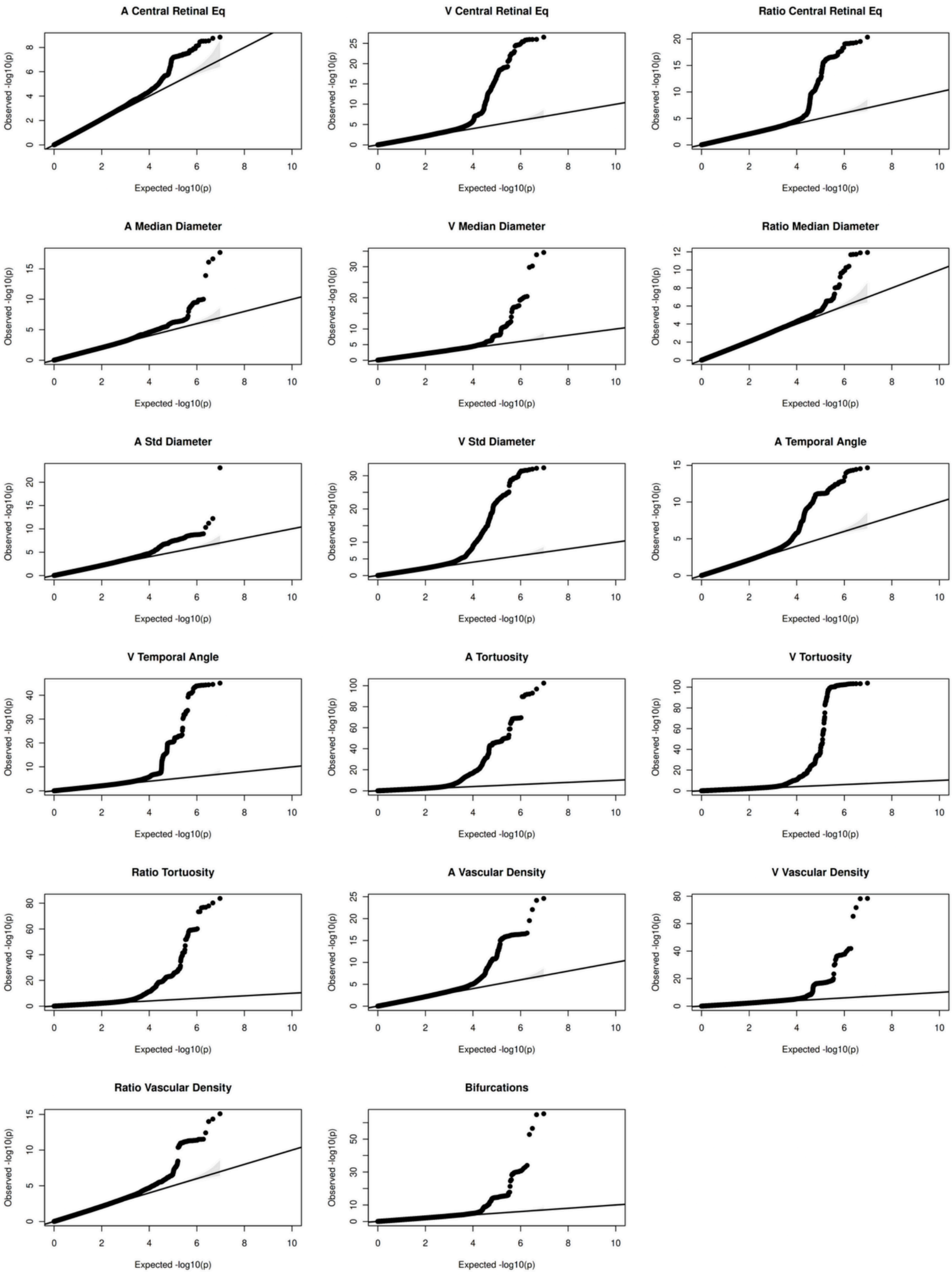


b) Quantile-Quantile (QQ) plots for all 17 retinal traits for Females.

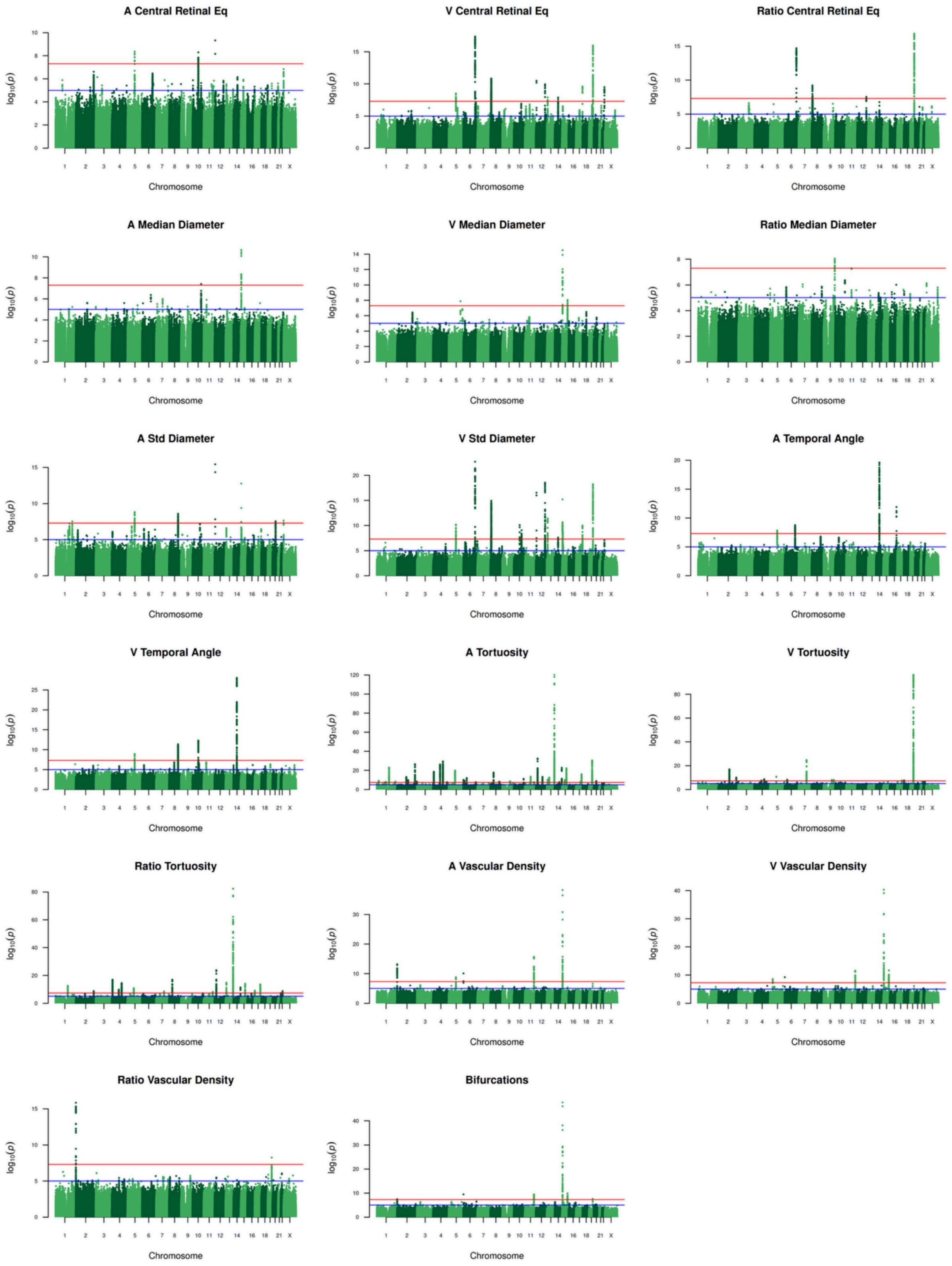


c) Manhattan plots of all 17 retinal traits for Males. Genome-wide significance (5e-8): red line, near genome-wide significance (1e-5): blue line.

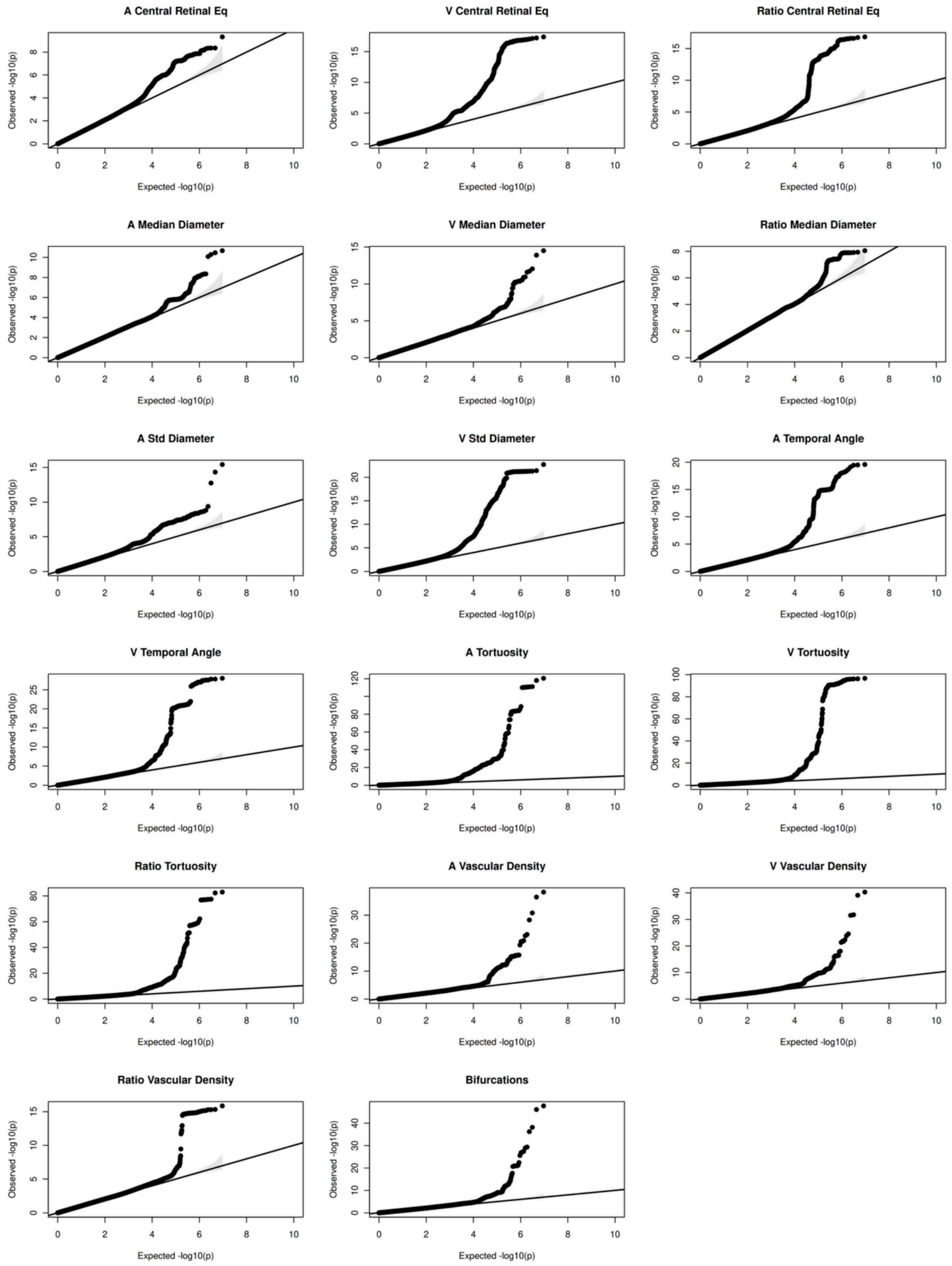


d) Quantile-Quantile (QQ) plots for all 17 retinal traits for Males.

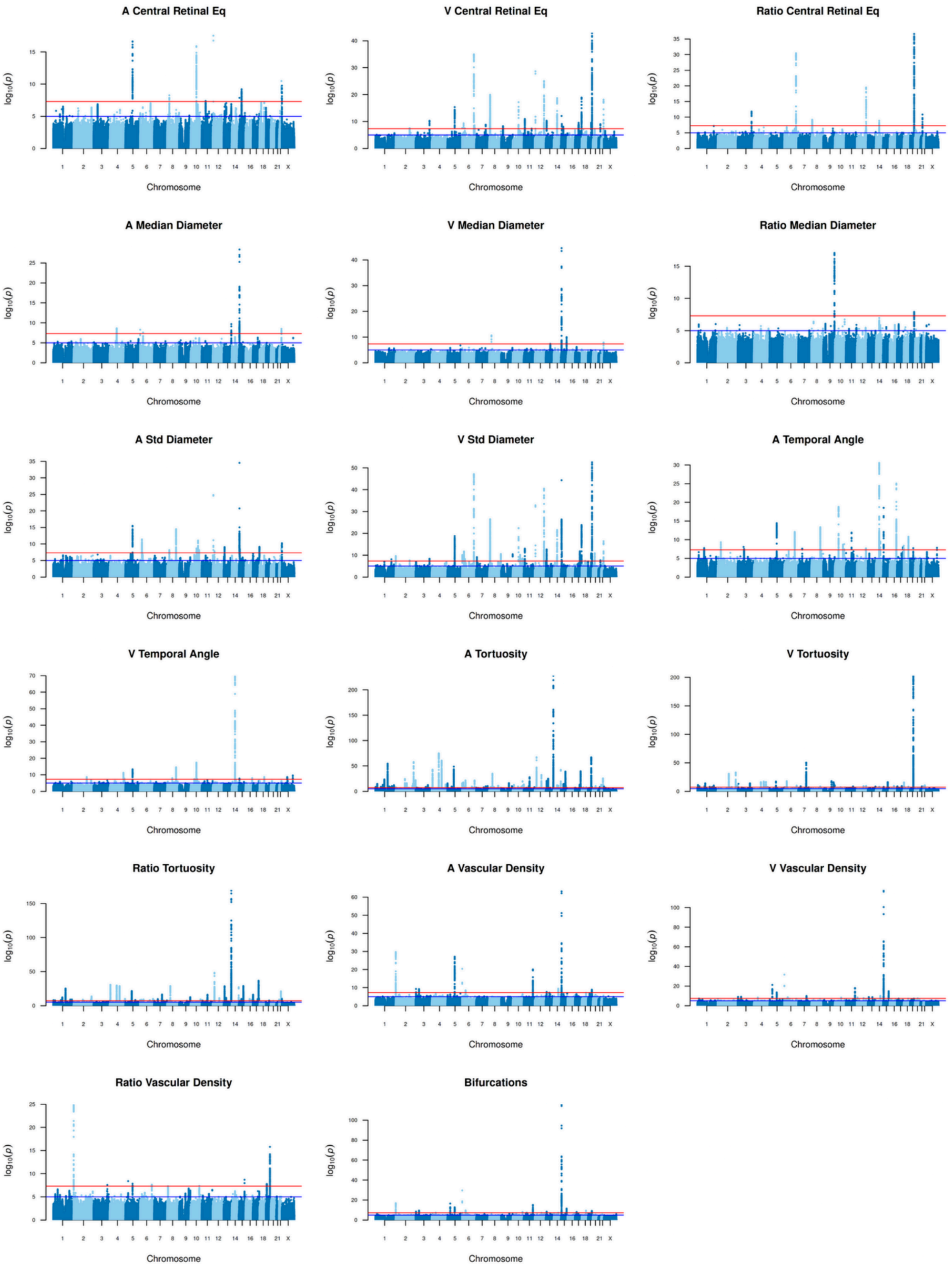


e) Manhattan plots of all 17 retinal traits for the combined sample. Genome-wide significance (5e-8): red line, near genome-wide significance (1e-5): blue line.

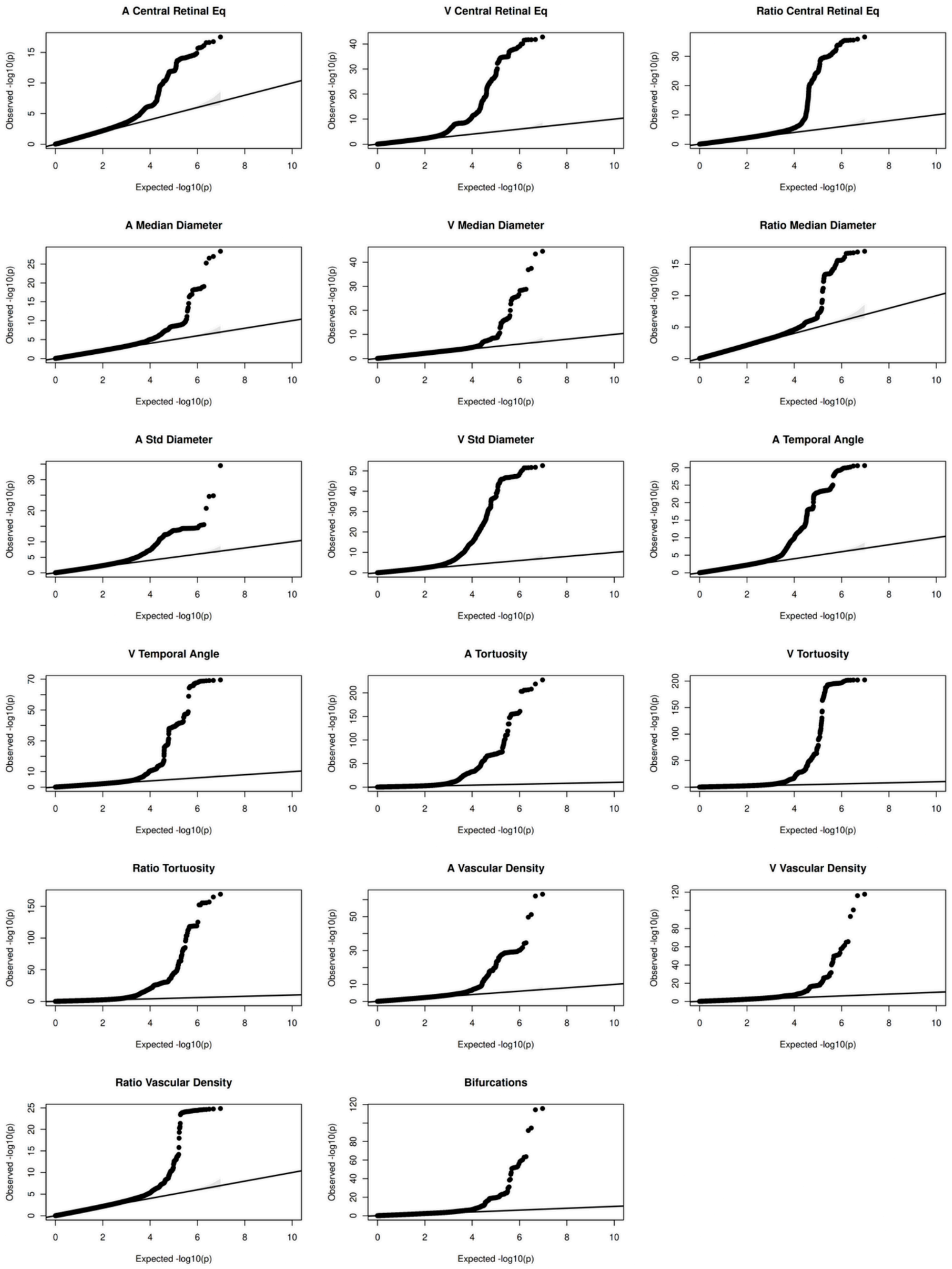


f) Quantile-Quantile (QQ) plots for all 17 retinal traits for the combined sample.

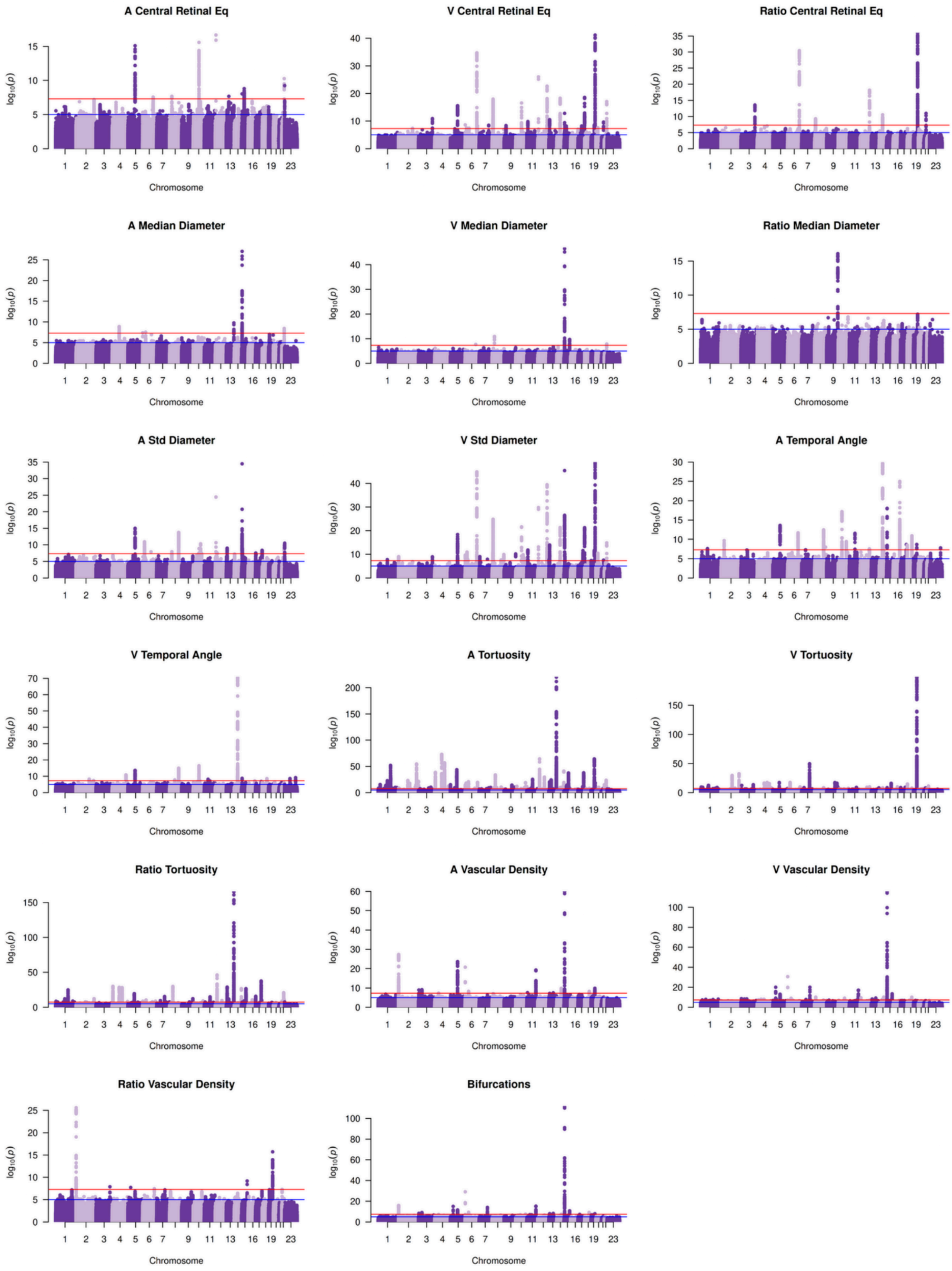


g) Manhattan plots of all 17 retinal traits for the meta-analysis. Genome-wide significance (5e-8): red line, near genome-wide significance (1e-5): blue line.

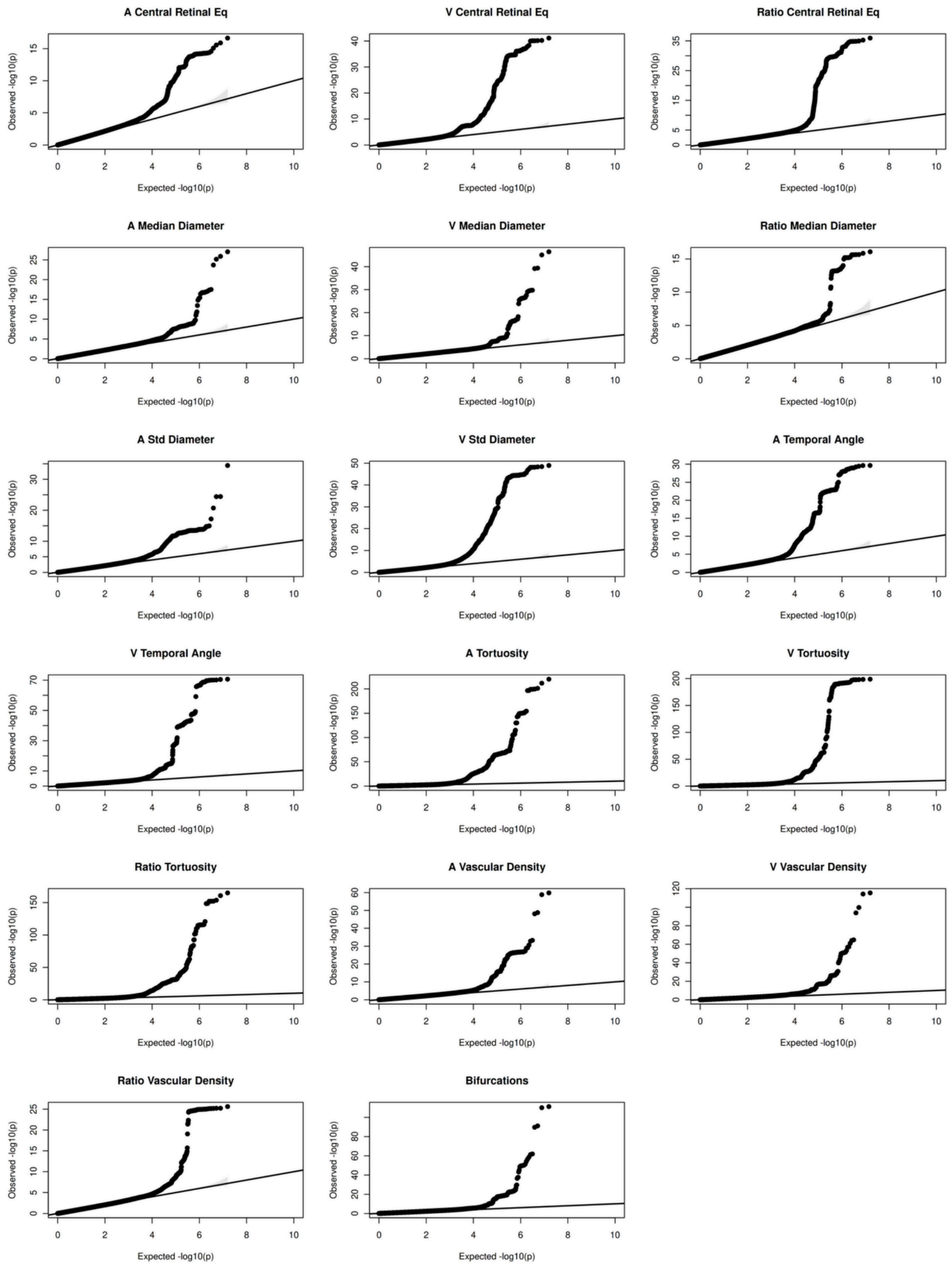


h) Quantile-Quantile (QQ) plots for all 17 retinal traits for the meta-analysis.

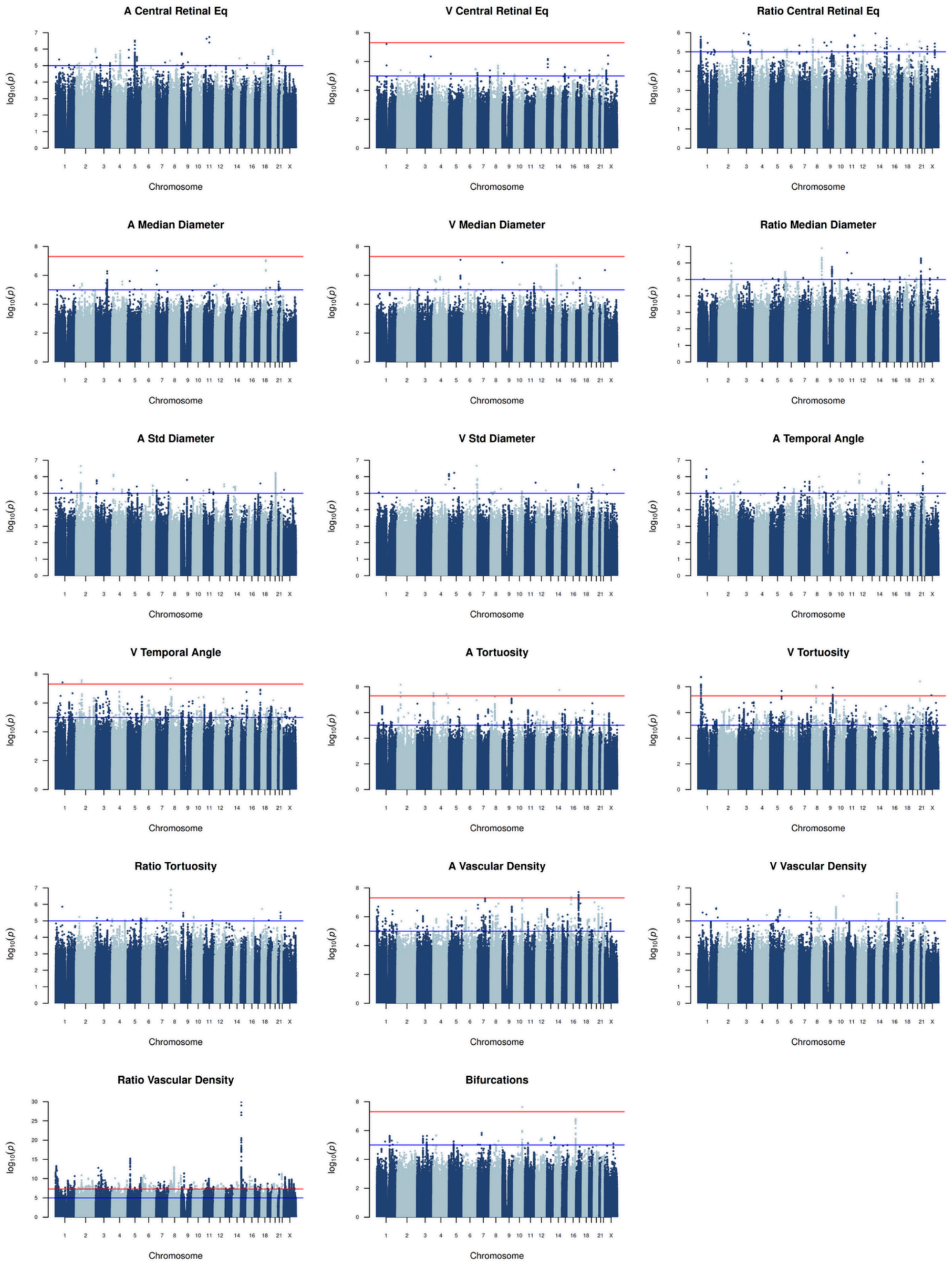


i) Manhattan plots of all 17 retinal traits for the interaction analysis. Genome-wide significance (5e-8): red line, near genome-wide significance (1e-5): blue line.

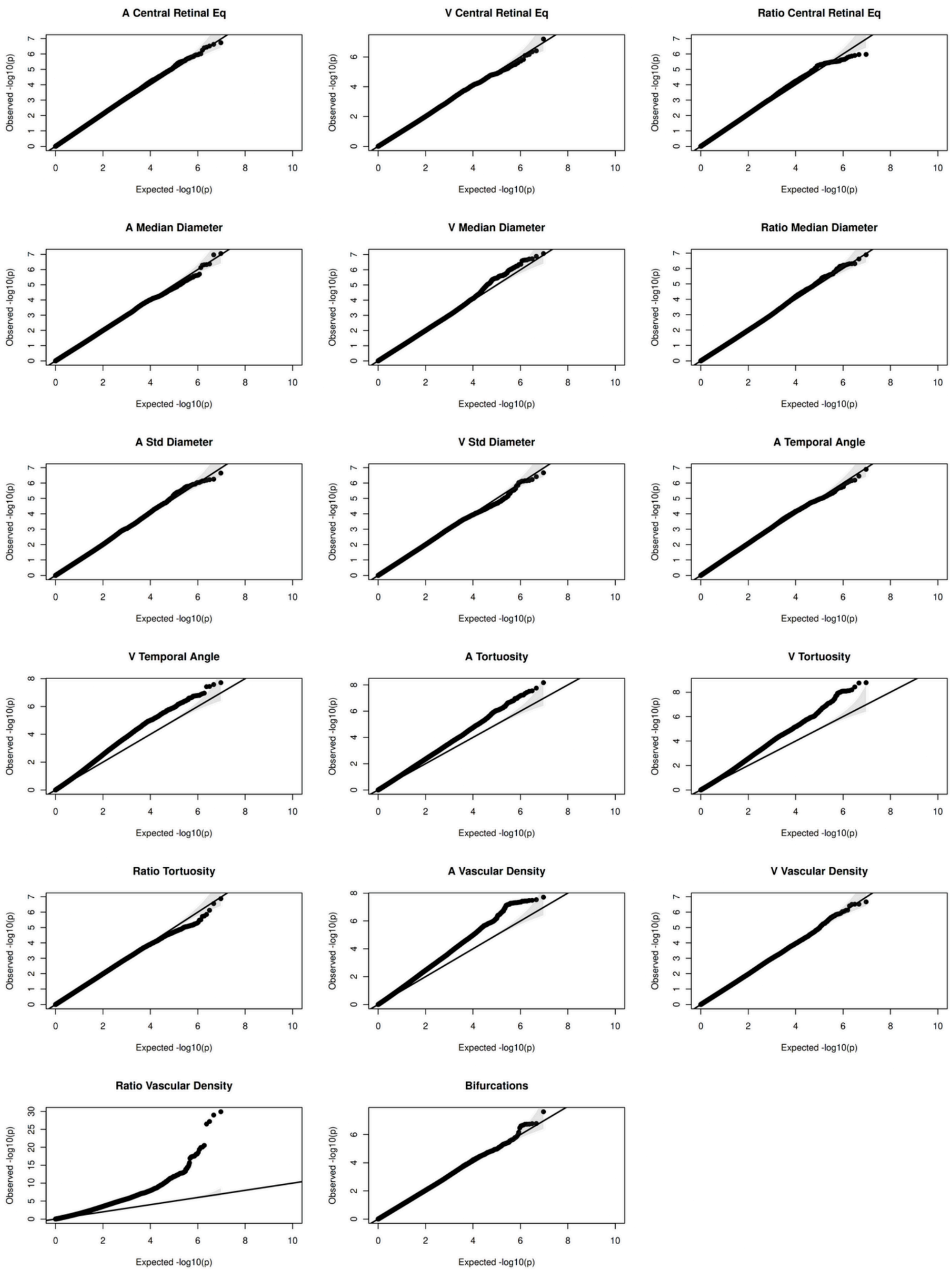


j) Quantile-Quantile (QQ) plots for all 17 retinal traits for the interaction analysis.

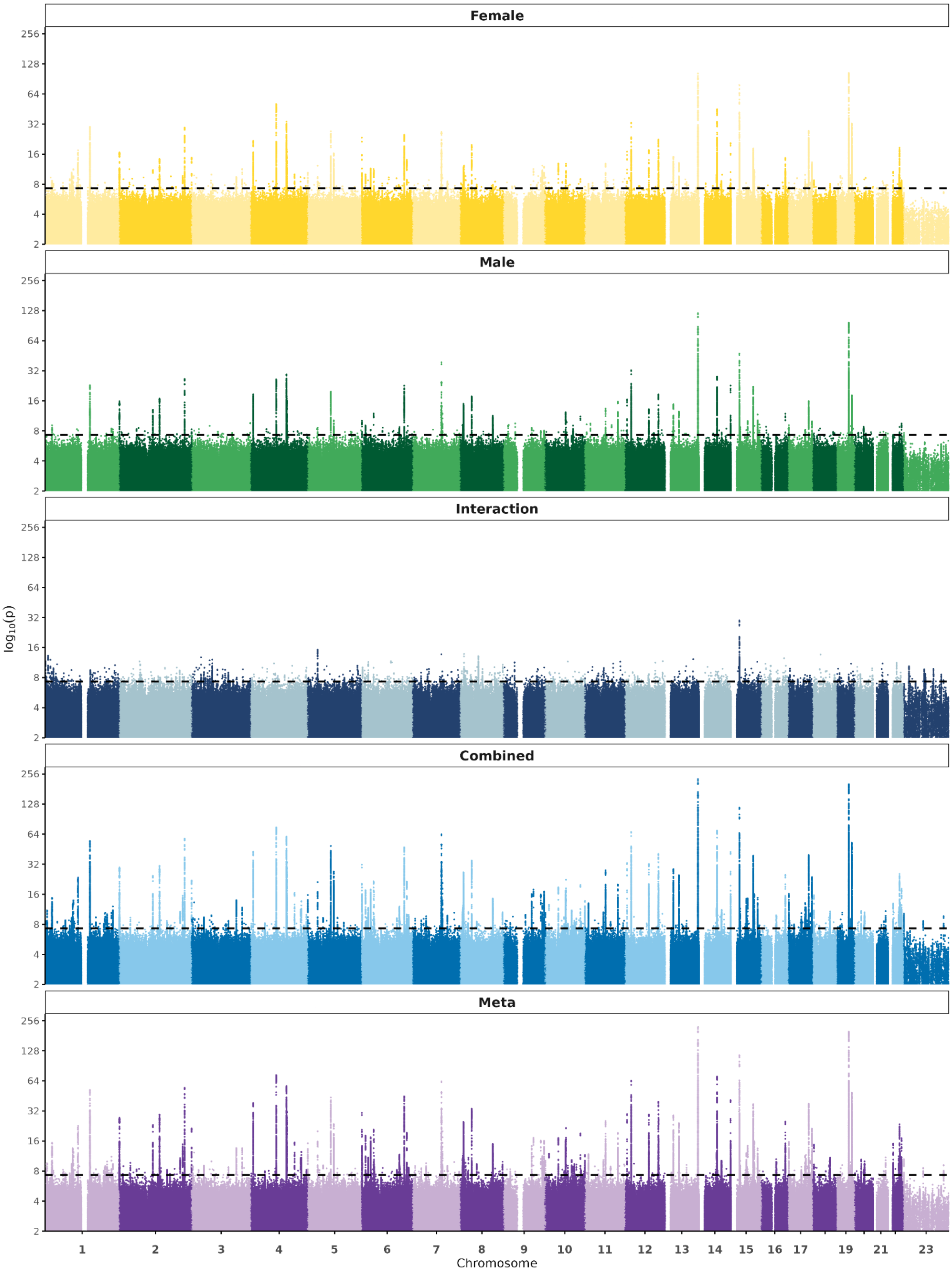


k) Overview. Thinned SNPs for all 17 retinal traits for all datasets/ analyses.

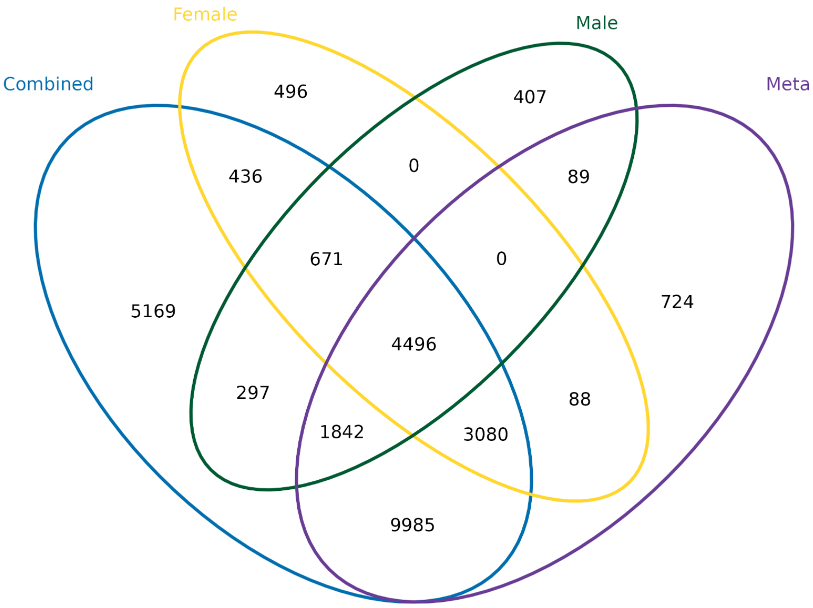


I) Venn Diagram of shared SNPs for all 17 IDPs for the combined, female, and male cohorts as well as the meta-analysis.

### Supplementary Figure 6: Trait-by-Genotype distribution

rs1129038 (Chr 15)

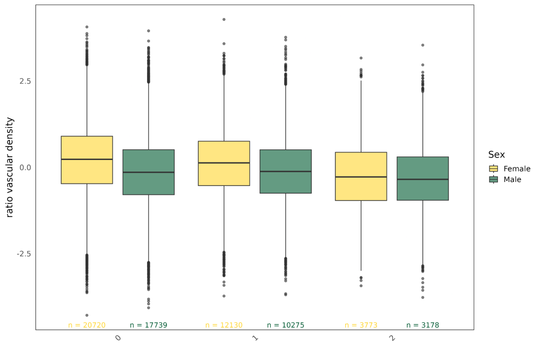


rs12898729 (Chr 15)

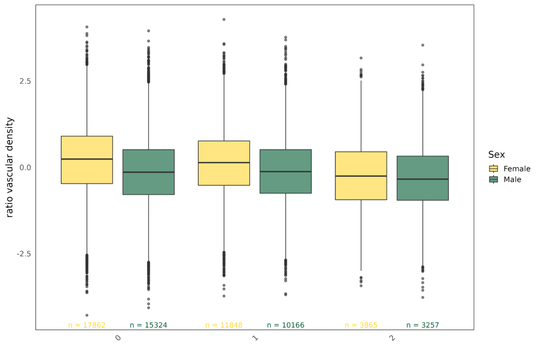


rs12912427 (Chr 15)

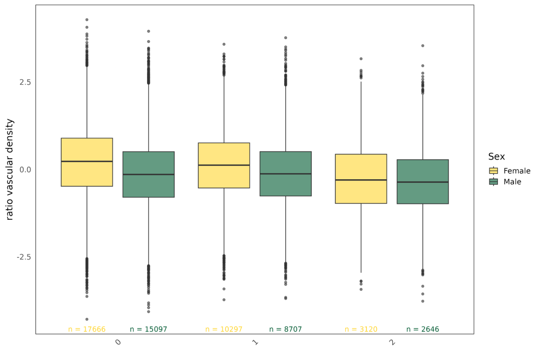


rs12913832 (Chr 15)

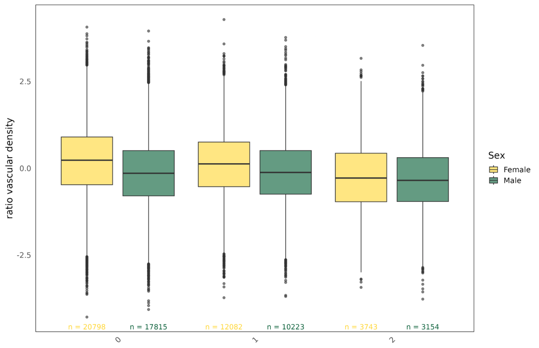


rs12916300 (Chr 15)

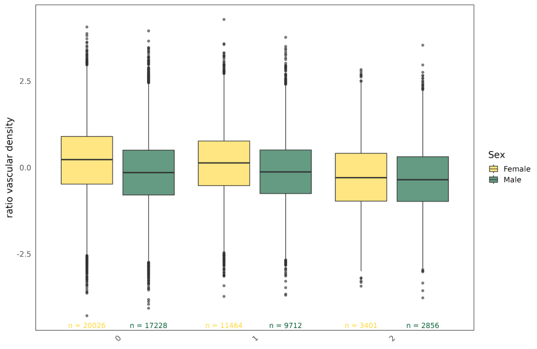


a) Boxplots showing trait per genotype distribution for females (yellow) and males (green). 0 = homozygous major, 1 = heterozygous, 2 = homozygous minor. Sample size per sex below each box.

rs1129038 ($\beta F$ = 0.069 , $\beta M$ = - 0.032)

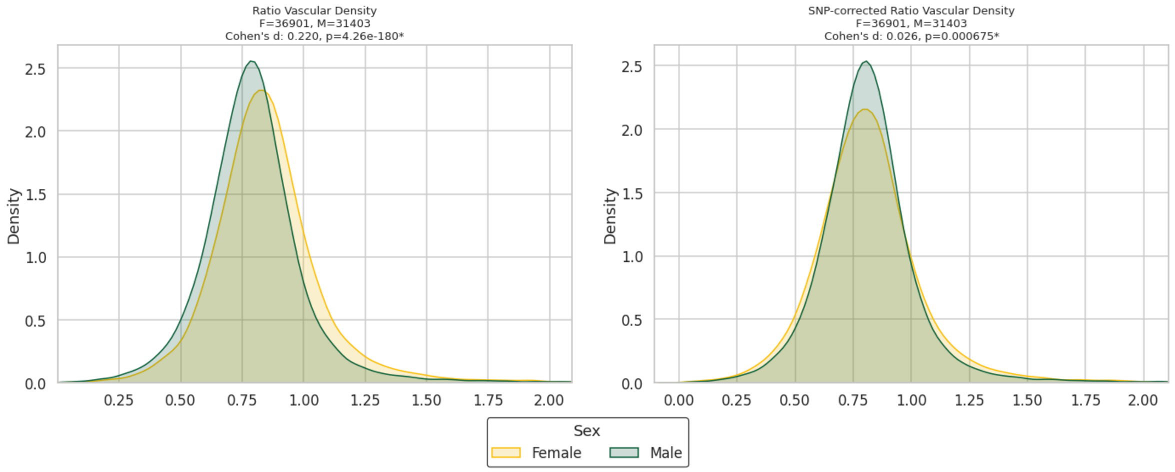


rs12898729 ($\beta F$ = 0.066 , $\beta M$ = - 0.026)

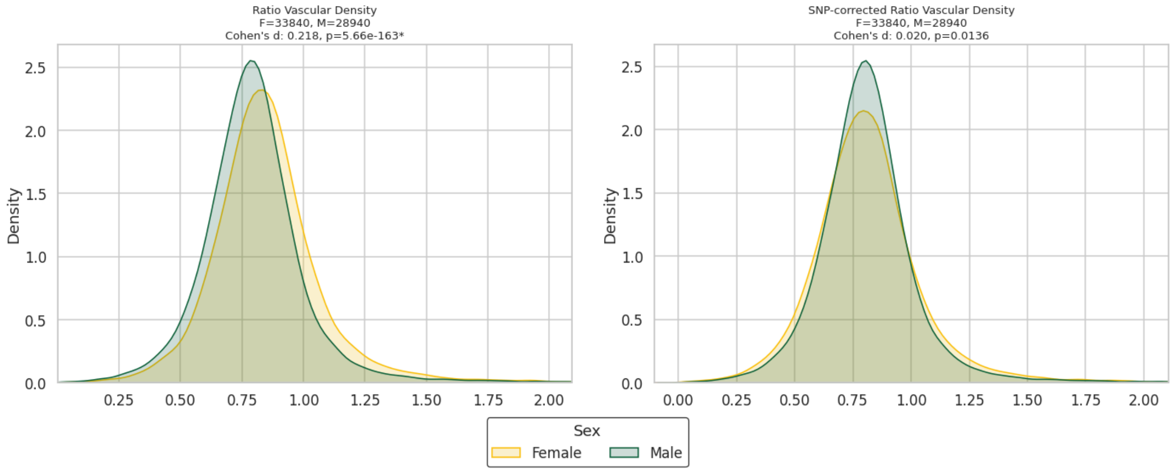


rs12912427 ($\beta F$ = 0.064 , $\beta M$ = - 0.036)

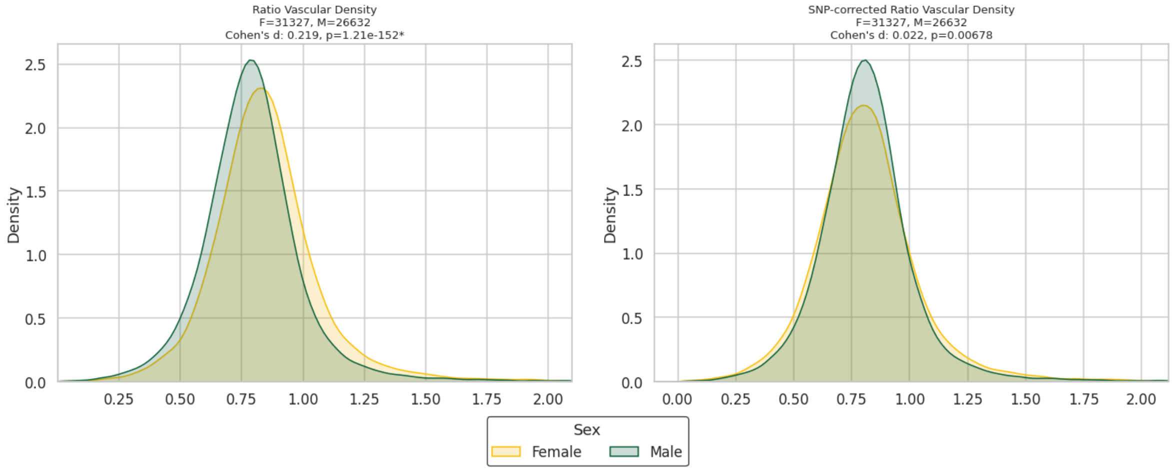

rs12913832 ($\beta F$ = 0.069 , $\beta M$ = - 0.032)

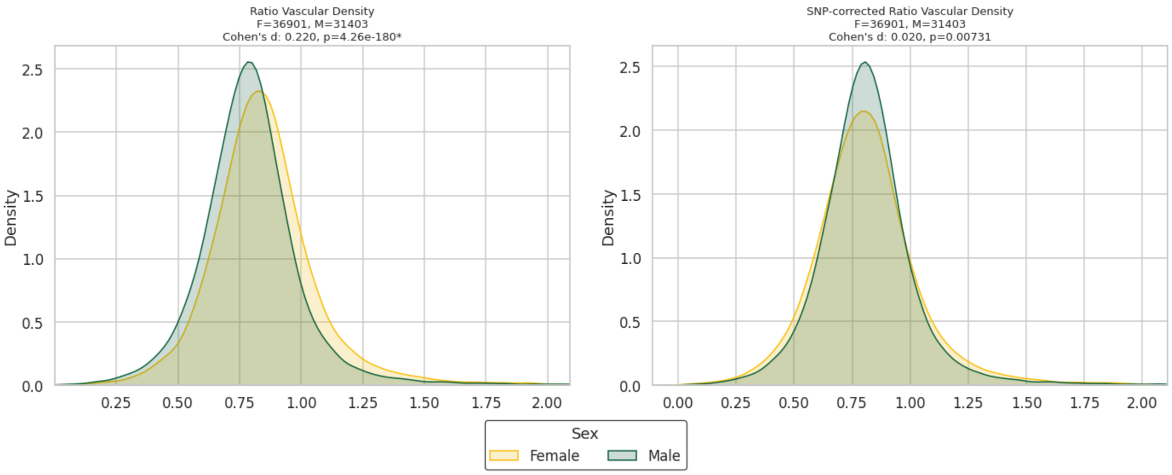


rs12916300 ($\beta F$ = 0.063 , $\beta M$ = - 0.031)

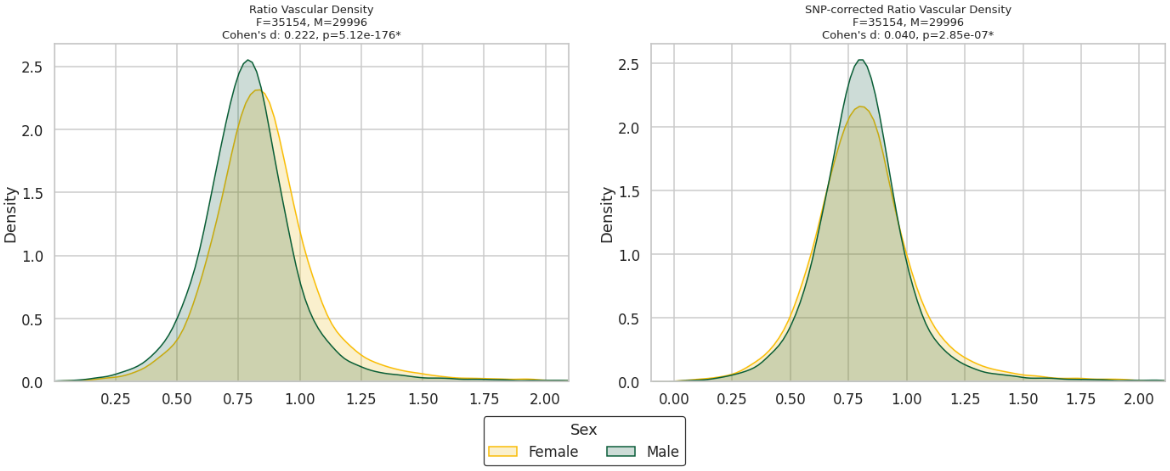


b) Comparison of raw A/V ratio vascular density (left) and SNP corrected A/V ratio vascular density (right).

### Supplementary Figures 7: Heritability results

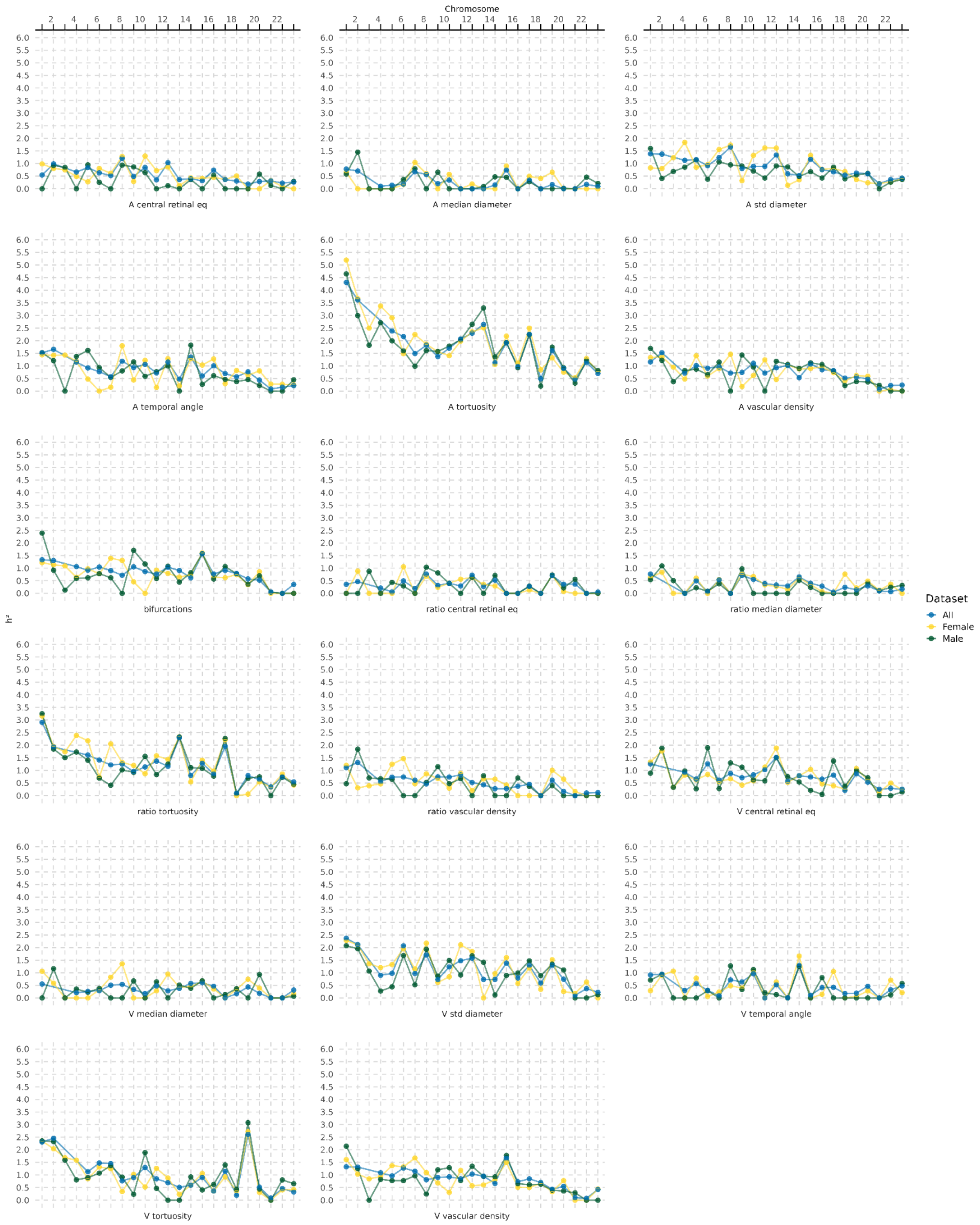


a) Trait and chromosome-wise SNP-heritability (in %), comparing the three datasets (Female, Male, Combined).

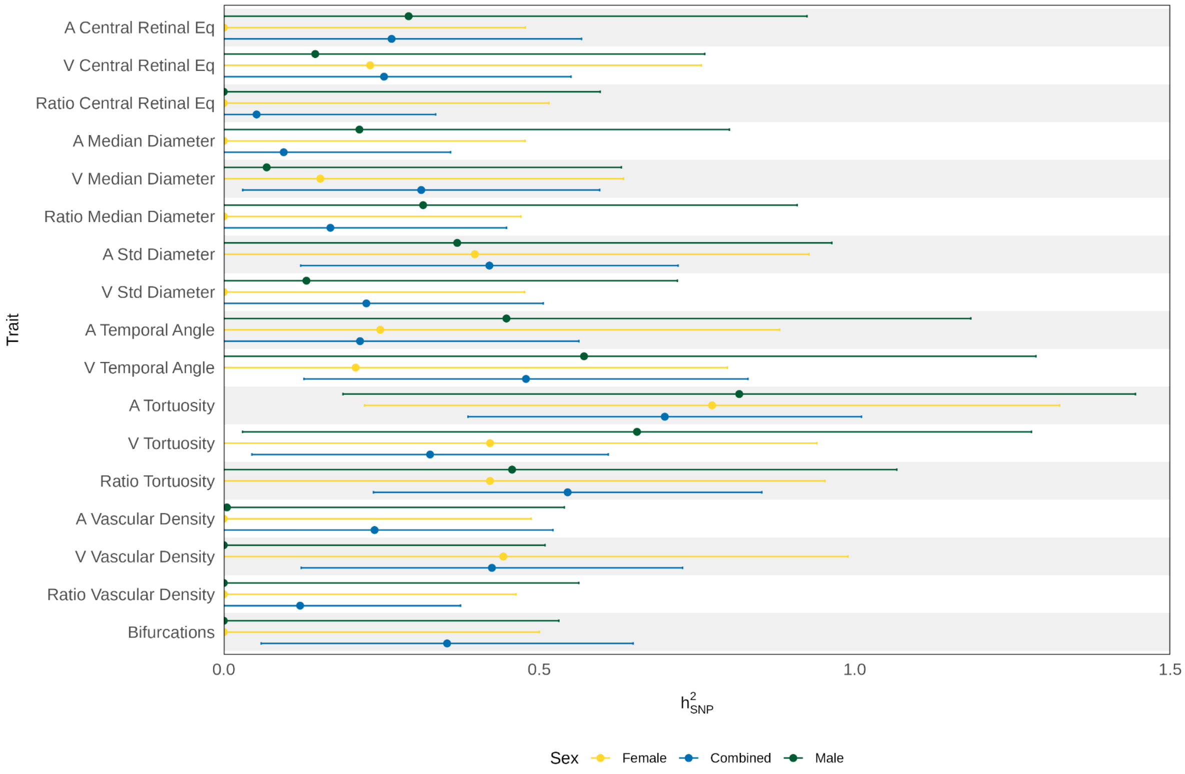


b) Zoomed in X-chromosomal SNP-heritability (in %) for all 17 retinal traits with 95%CIs.

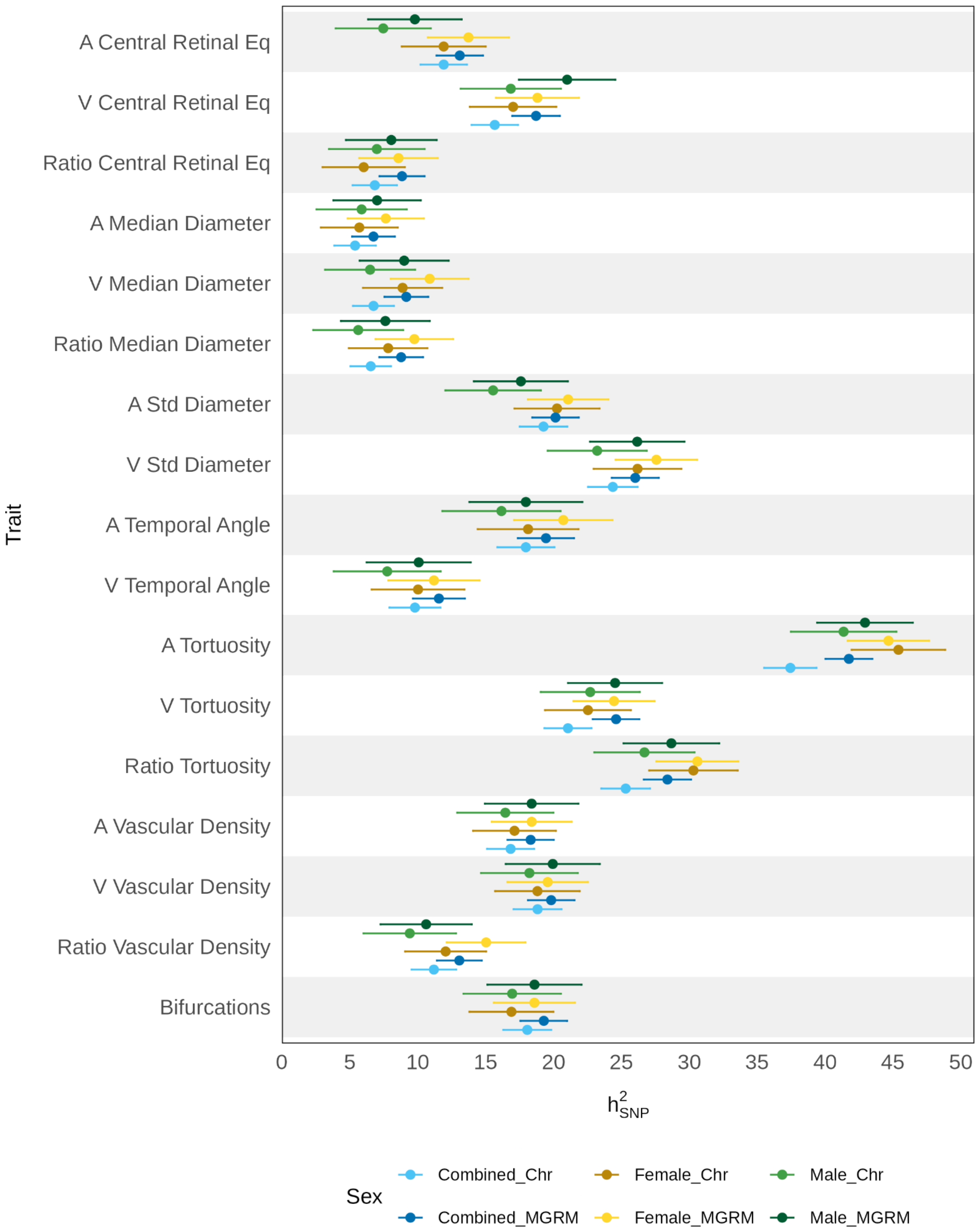


c) Comparison between additive model ("Chr") and Multi-component GREML ("MGRM") for all three cohorts. MGRM results presented in the main text.

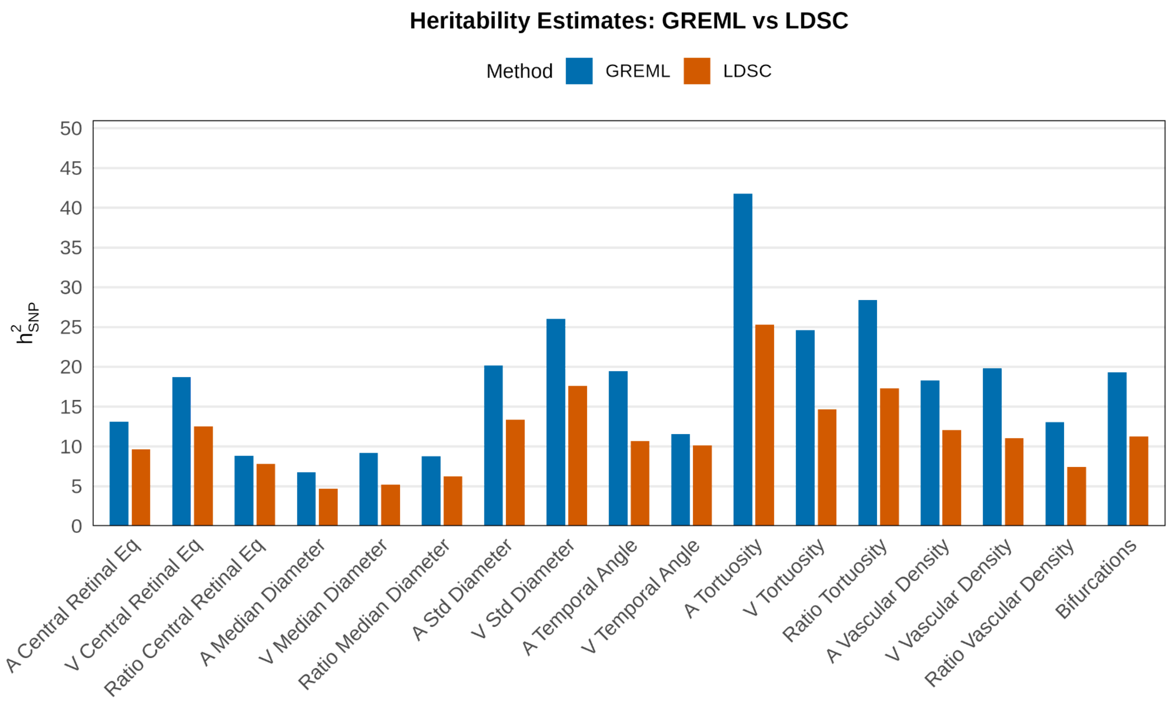


d) Comparison between LDSC estimates taken from Ortin Vela et al. (2024) (exact estimates not published but granted access to by the authors) and GREML estimates for the combined cohort.

### Supplementary Figure 8: Anti-Coherence Cross-Scoring Results

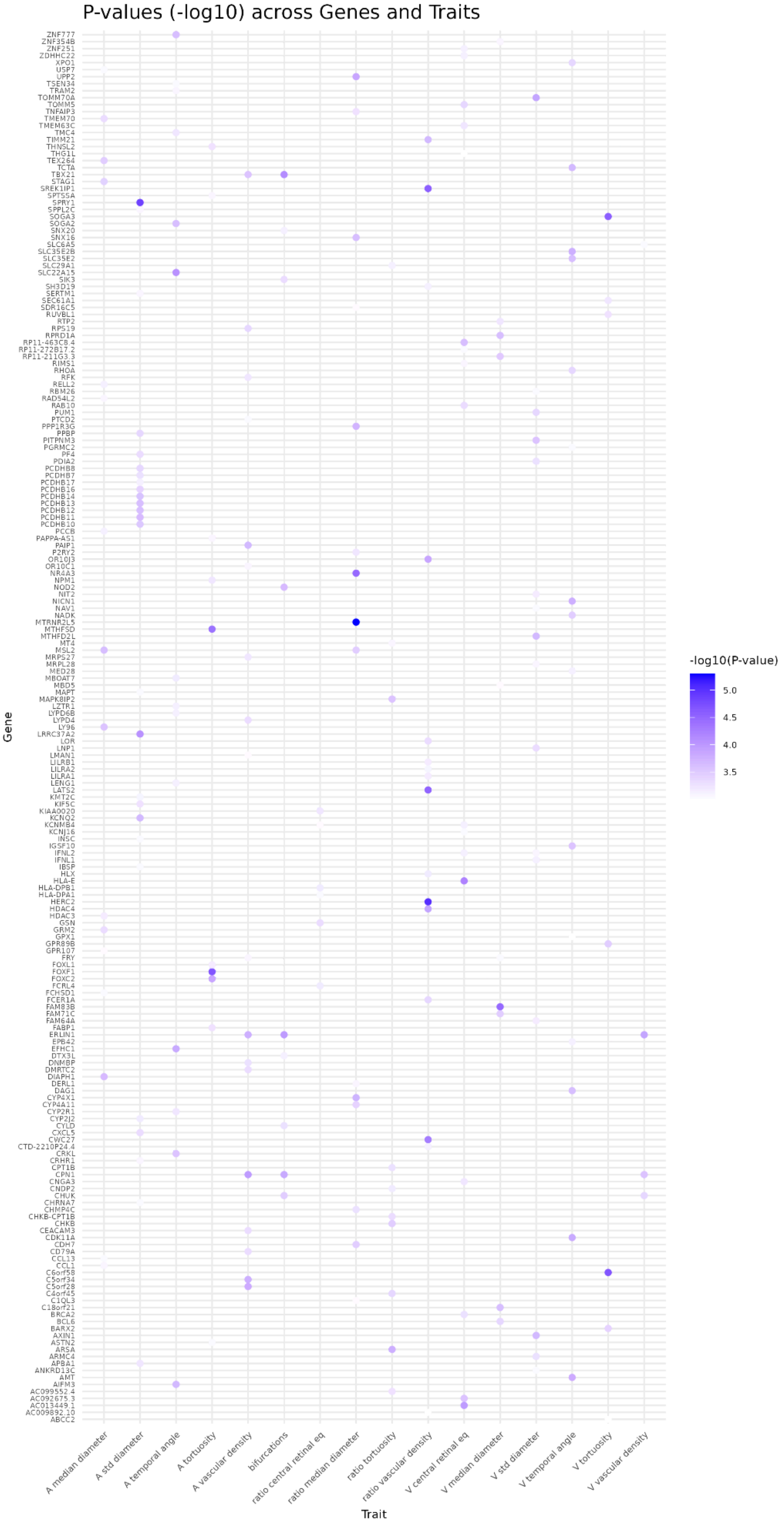


Results of PascalXs cross-scoring with non-significant anti-coherence.
