## Supplementary Information for "Sex-specific disease association and genetic architecture of retinal vascular traits"

### Supplementary Info 1: Distinction between sex as a biological variable and socio-cultural construct of gender

Whereas often used interchangeably or without definition, there is a clear distinction between sex and gender. This is crucial in biomedical research, with its inflation leading to imprecise scientific communication. “Sex” is a biological category normally assigned at birth (*sex assigned at birth, SAAB*) based on the observation of external genitalia or may be determined by the presence of a specific set of sex chromosomes, i.e., XX or XY. It is often treated as a binary variable, “female” or “male”, with exceptions occurring for intersex individuals. More recently, sex has been portrayed as a continuum as multiple factors, i.e., chromosomes, genes, and hormones, can lead to differential expression of sexual physical sex attributes[1]. Gender, on the other hand, is an umbrella term mainly used to refer to “gender identity”, defined as “a person’s deeply felt, inherent sense of being a boy, a man, or a male; a girl, a woman, or a female; or an alternative gender” (American Psychological Association, 2015) and is often deeply intertwined with “gender expression” or “gender roles/ stereotypes”, determined by the individual's socio-cultural context. Major health disparities have been recognised for individuals whose gender does not conform to their SAAB as well as those not falling into the binary sexual categories, generally referred to as sexual and gender minorities (Perry and LeBlanc, 2021).

[1] See this graphic developed in the course of a special issue on sex and gender at the ScientificAmerican: https://static.scientificamerican.com/sciam/cache/file/164FE5CE-FBA6-493F-B9EA84B04830354E_source.jpg,

### Supplementary Info 2: Covariates, Risk factors, diseases, and events

**Table 2.1 | Covariates used in the regression and genetic analyses.**

| Covariate | UKB Data Field | Field Name |
|---|---|---|
| Age | 21022 | Age at recruitment |
| Sex | 22001 | Genetic Sex |
| Assessment centre | 54 | UK Biobank assessment centre |
| Spherical power (left) | 5085 | Spherical power (left) |
| Spherical power (right) | 5084 | Spherical power (right) |
| Cylindrical power (left) | 5086 | Cylindrical power (left) |
| Cylindrical power (right) | 5087 | Cylindrical power (right) |
| Genotype batch | 22000 | Genotype measurement batch |
| Genetic PCs | 22009 | Genetic principal components |

**Table 2.2 | Covariates (Framingham risk factors) included in the survival analyses.**

| Risk factor | UKB Data Field | Field Name |
|---|---|---|
| Sex | 22001 | Genetic Sex |
| Systolic Blood Pressure | 4080 | Systolic blood pressure (automated reading) |
| Smoking status | 20116 | Smoking status |
| BMI | 21001 | Body Mass Index (BMI) |
| Total Cholesterol | 30690 | Cholesterol |
| HDL Cholesterol | 30760 | HDL cholesterol |
| Diabetes | 2976 | Age diabetes diagnosed |
| | 2443 | Diabetes diagnosed by doctor |
| | 30740 | Glucose |
| | 30750 | Glycated haemoglobin (HbA1c) |

| | | |
|---|---|---|
| Medicated for Hypertension | 6153 | Medication for cholesterol, blood pressure, diabetes, or take exogenous hormones |
| | 6177 | Medication for cholesterol, blood pressure or diabetes |
| Previous Cardiovascular Event | 131296, 131304, 131306, 131298, 131300, 131302, 42000, 42006, 42008, 42010, 131338, 131340, 131288, 131292 | See Table 3.2. |

**Table 2.3 | Diseases or risk factors used in the regression and survival analysis.**

| Disease/ Risk | UKB Data Field | Field Name |
|---|---|---|
| Systolic Blood Pressure | 4080 | Systolic blood pressure, automated reading |
| Diastolic Blood Pressure | 4079 | Diastolic blood pressure, automated reading |
| Hypertension | 6153 | Medication for cholesterol, blood pressure, diabetes, or take exogenous hormones |
| | 6177 | Medication for cholesterol, blood pressure or diabetes |
| | 4079 | Diastolic blood pressure, automated reading |
| | 4080 | Systolic blood pressure, automated reading |
| HDL | 23406 | HDL Cholesterol |
| LDL | 23405 | LDL Cholesterol |
| Triglycerides | 23407 | Total Triglycerides |
| Hba1c | 30750 | Glycated haemoglobin (HbA1c) |
| BMI | 21001 | Body Mass Index (BMI) |
| Pulse Rate | 102 | Pulse rate ,automated reading |
| Pulse Wave Arterial Stiffness Index | 21021 | Pulse Wave Arterial Stiffness Index |
| Alcohol Consumption | 1558 | Alcohol intake frequency |
| Smoking Pack-years | 20161 | Pack-years of smoking |
| Smoking Status | 20116 | Smoking status |
| Amblyopia | 6147 | Reason for glasses/contact lenses |

| | | |
|---|---|---|
| Presbyopia | 6147 | Reason for glasses/contact lenses |
| Myopia | 6147 | Reason for glasses/contact lenses |
| Astigmatism | 6147 | Reason for glasses/contact lenses |
| Hypermetropia | 6147 | Reason for glasses/contact lenses |
| Glaucoma | 6148 | Eye problems/disorders |
| Cataract | 6148 | Eye problems/disorders |
| Other Eye Condition | 6148 | Eye problems/disorders |
| Diabetes | – | see: Table 3.2. |
| Atherosclerosis | 131380 | Date I70 first reported (atherosclerosis) |
| IHD (Ischaemic Heart Disease) | 131296 | Date I20 first reported (angina pectoris) |
| | 131304 | Date I24 first reported (other acute ischaemic heart diseases) |
| | 131306 | Date I25 first reported (chronic ischaemic heart disease) |
| MI (Myocardial Infarction) | 131298 | Date I21 first reported (acute myocardial infarction) |
| | 131300 | Date I22 first reported (subsequent myocardial infarction) |
| | 131302 | Date I23 first reported (certain current complications following acute myocardial infarction) |
| | 42000 | Date of myocardial infarction |
| Stroke | 42006 | Date of stroke |
| | 42008 | Date of ischaemic stroke |
| | 42010 | Date of intracerebral haemorrhage |
| NIC (Non-Ischaemic Cardiomyopathies) | 131338 | Date I42 first reported (cardiomyopathy) |
| | 131340 | Date I43 first reported (cardiomyopathy in diseases classified elsewhere) |
| | 131288 | Date I11 first reported (hypertensive heart disease) |
| | 131292 | Date I13 first reported (hypertensive heart and renal disease) |
| Thrombotic Event | 131308 | Date I26 first reported (pulmonary embolism) |
| | 131388 | Date I74 first reported (arterial embolism and thrombosis) |
| | 131400 | Date I82 first reported (other venous embolism and thrombosis) |

| | | |
|---|---|---|
| Death | 40000 | Date of death |

**Table 2.4 | ICD10 codes used to determine cardiovascular-related (CVD) death.**

| ICD-10 Code | Description |
|---|---|
| I11.0 | Hypertensive heart disease with heart failure |
| I11.9 | Hypertensive heart disease without heart failure |
| I21.0 | Acute myocardial infarction |
| I21.9 | Acute myocardial infarction, unspecified |
| I25.1 | Atherosclerotic heart disease of native coronary artery |
| I25.3 | Aneurysm of heart |
| I25.9 | Chronic ischaemic heart disease |
| I26.9 | Pulmonary embolism without acute cor pulmonale |
| I48.9 | Atrial fibrillation and atrial flutter, unspecified |
| I49.9 | Cardiac arrhythmia, unspecified |
| I50.0 | Congestive heart failure |
| I60.8 | Other nontraumatic subarachnoid hemorrhage |
| I60.9 | Nontraumatic subarachnoid haemorrhage, unspecified |
| I61.8 | Other nontraumatic intracerebral hemorrhage |
| I61.9 | Nontraumatic intracerebral haemorrhage, unspecified |
| I62.9 | Nontraumatic intracranial haemorrhage, unspecified |
| I63.2 | Cerebral infarction due to unspecified occlusion or stenosis of precerebral arteries |
| I63.3 | Cerebral infarction due to thrombosis of cerebral arteries |
| I63.9 | Cerebral infarction, unspecified |
| I64 | Stroke, not specified as haemorrhage or infarction. |
| I80.2 | Phlebitis and thrombophlebitis of other deep vessels of lower extremities |
| I81 | Portal vein thrombosis |

**Table 2.5 | Sample size, events, and follow-up for the survival analysis endpoints.** Characteristics of the survival analysis cohorts for different endpoints after excluding missing covariates.

| Endpoint | Characteristics | Overall cohort (N = 58608) | Females cohort (N = 31422) | Males cohort (N = 27186) |
|---|---|---|---|---|
| Death | Events | 4539 (7.75%) | 1855 (5.90%) | 2684 (9.87%) |
| | Follow-up [years] | 14.62 | 14.62 | 14.61 |
| CVD Death | Events | 689 (1.18%) | 236 (0.07%) | 453 (1.67%) |
| | Follow-up [years] | 14.64 | 14.64 | 14.63 |
| CVD Event | Events | 7006 (11.95%) | 2656 (8.45%) | 4350 (16%) |
| | Follow-up [years] | 14.61 | 14.62 | 14.60 |
| IHD Event | Events | 4670 (7.97%) | 1653 (5.26%) | 3017 (11.10%) |
| | Follow-up [years] | 14.62 | 14.63 | 14.61 |
| MI Event | Events | 1743 (2.97%) | 545 (1.73%) | 1198 (4.41%) |
| | Follow-up [years] | 14.63 | | |
| Stroke Event | Events | 1291 (2.20%) | 550 (1.75%) | 741 (2.73%) |
| | Follow-up [years] | 14.63 | | |

**Supplementary Info 3: Linkage Disequilibrium (LD) for SNPs on Chr. 15**

We estimated LD between the SNPs on chromosome 15, significant in the individual female GWAS for ratio vascular density, which further showed significant differences in beta coefficients to the male GWAS using Ensembls Linkage Disequilibrium Calculator (https://www.ensembl.org/Homo_sapiens/Tools/LD/) in the 1000 Genomes, GBR population *(Population 1000GENOMES:phase_3:GBR).*

| Variant 1 | V1: consequence | Variant 2 | V2: consequence | $r^2$ | D’ |
|---|---|---|---|---|---|
| rs12898729 | intron variant | rs12912427 | intron variant | 0.645 | 0.956 |
| rs12916300 | intron variant | rs12912427 | intron variant | 0.754 | 0.919 |
| rs1129038 | 3’ UTR variant | rs12898729 | intron variant | 0.762 | 1 |
| rs12913832 | intron variant | rs12898729 | intron variant | 0.762 | 1 |
| rs12898729 | intron variant | rs12916300 | intron variant | 0.790 | 1 |
| rs1129038 | 3’ UTR variant | rs12912427 | intron variant | 0.854 | 0.960 |
| rs12913832 | intron variant | rs12912427 | intron variant | 0.854 | 0.960 |
| rs1129038 | 3’ UTR variant | rs12916300 | intron variant | 0.893 | 0.962 |
| rs12913832 | intron variant | rs12916300 | intron variant | 0.893 | 0.962 |
| rs1129038 | 3’ UTR variant | rs12913832 | intron variant | 1 | 1 |

*r2 and D’ values rounded to third decimal.

**Supplementary Info 4: Trait-wise heritability per cohort**

| Trait | Cohort | $h^2_{SNP}$ | $h^2_{SNP}$ SE |
|---|---|---|---|
| A Central Retinal Eq | Female | 0.137 | 0.015 |
| | Male | 0.098 | 0.018 |
| | Combined | 0.131 | 0.009 |
| V Central Retinal Eq | Female | 0.188 | 0.016 |
| | Male | 0.210 | 0.018 |
| | Combined | 0.187 | 0.009 |
| Ratio Central Retinal Eq | Female | 0.086 | 0.015 |
| | Male | 0.081 | 0.017 |
| | Combined | 0.088 | 0.008 |
| A Median Diameter | Female | 0.076 | 0.014 |
| | Male | 0.070 | 0.016 |
| | Combined | 0.067 | 0.008 |
| V Median Diameter | Female | 0.109 | 0.015 |
| | Male | 0.090 | 0.017 |
| | Combined | 0.092 | 0.008 |
| Ratio Median Diameter | Female | 0.098 | 0.015 |
| | Male | 0.076 | 0.017 |
| | Combined | 0.088 | 0.008 |
| A Std Diameter | Female | 0.211 | 0.015 |
| | Male | 0.176 | 0.018 |
| | Combined | 0.202 | 0.009 |
| V Std Diameter | Female | 0.276 | 0.015 |
| | Male | 0.262 | 0.018 |
| | Combined | 0.260 | 0.009 |
| A Temporal Angle | Female | 0.207 | 0.019 |
| | Male | 0.180 | 0.021 |
| | Combined | 0.194 | 0.011 |

| | | | |
|---|---|---|---|
| V Temporal Angle | Female | 0.112 | 0.017 |
| | Male | 0.101 | 0.020 |
| | Combined | 0.116 | 0.010 |
| A Tortuosity | Female | 0.447 | 0.015 |
| | Male | 0.429 | 0.018 |
| | Combined | 0.418 | 0.009 |
| V Tortuosity | Female | 0.245 | 0.015 |
| | Male | 0.245 | 0.018 |
| | Combined | 0.246 | 0.009 |
| ratio Tortuosity | Female | 0.306 | 0.015 |
| | Male | 0.287 | 0.018 |
| | Combined | 0.284 | 0.009 |
| A Vascular Density | Female | 0.184 | 0.015 |
| | Male | 0.184 | 0.018 |
| | Combined | 0.183 | 0.009 |
| V Vascular Density | Female | 0.196 | 0.015 |
| | Male | 0.199 | 0.018 |
| | Combined | 0.198 | 0.009 |
| ratio Vascular Density | Female | 0.150 | 0.015 |
| | Male | 0.106 | 0.017 |
| | Combined | 0.131 | 0.008 |
| Bifurcations | Female | 0.186 | 0.015 |
| | Male | 0.186 | 0.018 |
| | Combined | 0.193 | 0.009 |
